# Enzyme family-centred approach identifies helicases as recurrent hemizygous tumour suppressor genes

**DOI:** 10.1101/2024.08.13.607019

**Authors:** Karolin Voßgröne, Francesco Favero, Krushanka Kashyap, F. Germán Rodríguez-González, André Vidas Olsen, Xin Li, Balca R. Mardin, Joachim Weischenfeldt, Claus S. Sørensen

**Affiliations:** Biotech Research and Innovation Centre, University of Copenhagen, Ole Maaløes Vej 5, Copenhagen 2200 N, Denmark; Finsen Laboratory, Copenhagen University Hospital - Rigshospitalet, Copenhagen, Denmark; BioMed X Institute (GmbH), Heidelberg, Germany; Charité - Universitätsmedizin Berlin, Berlin, Germany

**Author notes:** First authors (Karolin Voßgröne, Francesco Favero).

## Abstract

An important goal in cancer research is to identify driver genes and mutations. Reasoning that such mutations often alter enzymatic functions, we investigated the cancer driver role of enzyme families. Using pan-cancer genomic data and established driver mutation catalogues, we found an unexpectedly high rate of mutations in helicases, making helicases the most frequently mutated enzyme family in cancer. Based on both functional perturbation screens and cancer genomic analyses, we provide evidence that cancers with mutated helicases converge on increased genomic instability and faulty DNA repair. We identify a striking phenotype in cells with loss of the helicase Aquarius (AQR). AQR was exclusively hemizygous lost in cancer genomes, which was associated with elevated levels of structural variants and point mutation signatures indicative of homologous recombination deficiency. Finally, we leverage large dependency maps to show that hemizygous loss is a common tumour suppression mechanism among helicases. In summary, we uncover a striking frequency of mutated helicases with key roles in genomic maintenance, and we nominate novel hemizygous cancer driver genes including AQR.

## Introduction

Cancer is fuelled by driver mutations in genes that promote and orchestrate cancer development and progression. The mutated oncogenes and tumour suppressor genes (TSGs) have classically been identified based on the recurrence of aberrations affecting the genes, with gain-of-function (GoF) mutations of oncogenes and loss-of-function (LoF) mutations of TSGs. Cancer drivers have been challenging to identify due to their context-dependence, both in terms of tissue type, cell-of-origin and dependence on dysregulated pathways and other cancer genes. Statistical concepts and approaches have been developed to identify recurrently mutated genes within and among different cancer types^1,2^ and enrichment of mutations in specific signalling pathways^3,4^. Despite advances, genetic alterations contributing to tumour development and progression are not well-characterised for a substantial number of cancers. Current approaches seek to identify genes or loci that are more frequently mutated than expected by chance but do generally not consider *a priori* the function or activity of the affected genes.

An orthogonal approach is to harness information on the molecular function and activity of the mutated protein. A key function is enzymatic activity, such as phosphorylation and dephosphorylation of proteins by kinases and phosphatases, respectively. Enzymatic activities catalyse thousands of biochemical reactions in the cell^5^. Specifically, biochemical classification of enzymes is based on common biochemical activity rather than sequence similarity. Accordingly, sequences within a class are often highly divergent, thus, sequence-related enzyme groups within a class are termed families^6^. A number of enzyme families are associated with many cancer drivers such as kinases and phosphatases^7,8^, however, it remains to be determined if additional enzyme families aggregate cancer drivers and if this could be harnessed further.

To address this gap in cancer knowledge, we develop a new enzyme-family driver identification approach that combines information of enzyme-wide molecular activities and mutational frequencies. This pinpoints cancer driver functions towards enzyme types and prompts unbiased genotype-phenotype categorization and characterization. Here, we carried out an enzyme family-centric approach based on genomic data from 5,844 cancer patients. We identify an unexpectedly high rate of mutations across various cancer types in helicases and nucleases, enzymes involved in unwinding and processing of nucleic acids. Using a high-throughput functional phenotype screening approach, we uncover a major biological role for helicases in genome maintenance. Among this family of enzymes, we find and functionally characterise a particular striking phenotype for *AQR* and demonstrate that hemizygous loss of AQR is associated with distinct structural variant and mutation signatures in cancer genomes. In summary, the enzyme-family approach uncovers helicases as commonly mutated in cancer, point to a hemizygous driver mechanism particularly abundant in helicases, and a putative therapeutic option in tumours with hemizygous AQR loss.

## Results

### Pan-cancer mutation enrichment analysis reveals helicases as a recurrently mutated enzyme family

To enable a comprehensive analysis of mutations in enzyme families, we first identified genes belonging to the seven main enzyme classes: oxidoreductases, transferases, hydrolases, lyases, isomerases, ligases and translocases ^9^. We analysed the pan-cancer mutation burden for all genes in each enzyme class using publicly available genomic data generated by the TCGA Research Network (https://www.cancer.gov/tcga) and METABRIC^10^ across 18 different cancer types. By comparing the observed number of mutations in each class to the expected distribution of mutations, we found an increased mutational frequency in translocases, hydrolases and transferases (Effect size=6.8, 6.3, 6.7, *P*=7.7x10^-39^, 1.5x10x^-^ ^37^, 2.3x10^-36^, respectively, Fisher’s combined probability, Fig. 1A). This prompted us to investigate the associated enzyme families, a further division of the main enzyme classes characterized by amino acid sequence and structural similarities. To this end, we identified genes belonging to 11 enzyme families including 409 oxidoreductases, 25 glycosidases, 159 G proteins, 365 kinases, 75 lipases, 73 methyltransferases, 173 phosphatases, 267 proteases, 18 transaminases, 162 helicases and 156 nucleases^11^. Oxygenases were removed from further analyses due to a high redundancy with oxidoreductase enzyme family and a low number of unique enzymes. Adjusting for the number and size of genes in each family, we found cancer-related kinase, protease and phosphatase enzyme families to be significantly mutated across all cancer types (*P* = 1.9x10^-30^, 6x10^-21^, 1.1x10^-15^ respectively, 10,000 permutations, Fisher’s combined probability). Surprisingly, we found helicases to be the most frequently mutated enzyme family (effect size 7.4, *P* = 7.1x10^-45^, Fisher’s combined probability, Fig. 1A-B). At the individual gene level, we found cancer associated genes such as *ATRX*, *POLQ* and *SMARCA4* to be frequently mutated helicases across different cancers (Fig. 1C).

**Fig. 1.**
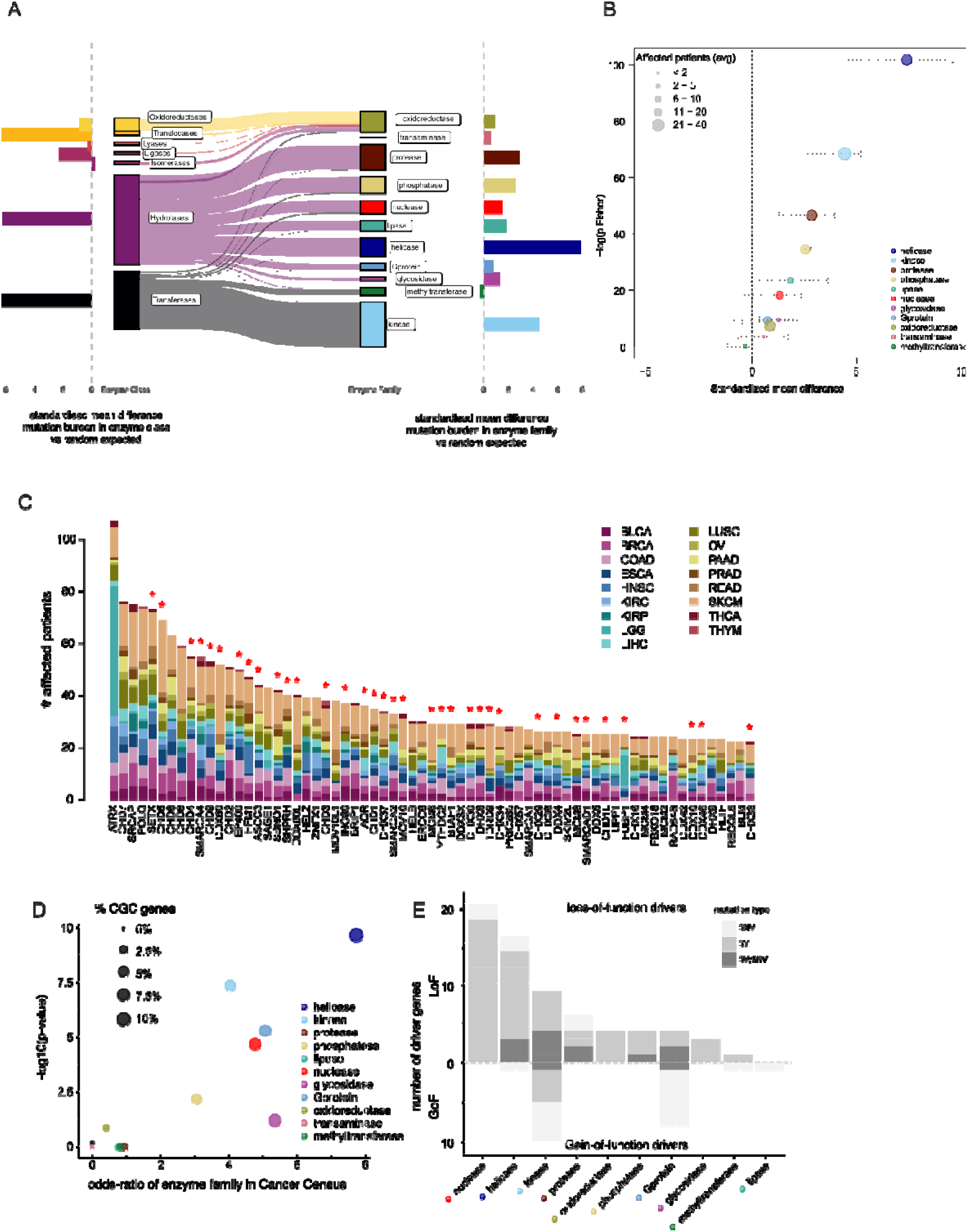
Cancer genomic analyses identify helicases as a recurrently mutated enzyme family. A) Enzyme classes partitioned as UniProtKB first-level annotation (left) and PantherDB annotation for selected enzyme families (right), The horizontal bar plots on the left and right sides correspond to the average effect size. B) Pan-cancer mutation enrichment significance by enzyme families, the y-axis corresponds to the Fisher combined p-value (-log) of 17 cancer studies from TCGA and METABRIC, the x-axis corresponds to the average effect size (coloured points) computed as the mean difference of each enzyme family to a permuted background set. The range of effect size across all studied cancer types is represented as horizontal dotted lines. C) Occurrence of somatic mutations in helicases: the most frequently mutated helicase genes ordered by occurrence. Each column corresponds to a gene, the colours in each column identify different tumour types, the height of a column corresponds to the number of patients across 17 different tumour type studies (legend) carrying a mutation in the corresponding gene. Red asterisk on top of the bars denotes a significant GISTIC copy number loss peak. D) Driver genes enrichment analysis for the 11 enzyme families. Cancer Gene Census (CGC) catalogue from COSMIC (version 95) was used to test significant enrichment for known driver genes in each enzyme family. The significance level (-log P-value, x-axis) versus enrichment of CGC genes in each enzyme family (odds-ratio, x-axis). The dot size is proportional to the fraction of genes in the enzyme family present in the CGC. E) For each enzyme family (x-axis), the number of CGC driver genes classified as LoF (y-axis, top barplot) and GoF (y-axis, bottom barplot) separated by the type of driver mutation involving SNVs (light grey), SVs (medium grey) or both SNVs and SVs (dark grey).

To explore whether and to what extent these enzyme families were enriched for driver genes, we next performed an unbiased enrichment analysis using the COSMIC Cancer Gene Census (CGC) catalogue of known driver genes (version 95). In agreement with our pan-cancer mutation burden analysis, helicases showed the strongest enrichment among CGC genes (odds-ratio 7.7, *P*<1x10^-10^, Fisher’s exact test, FDR-corrected p-value, Fig. 1D), with 18 out of 145 helicases listed as CGC, followed by kinases, phosphatases, and G proteins. Helicase CGC genes were associated with escaping programmed cell death and genome instability (Extended data Fig. 1A and Supplementary Table 1).

To gain more insights into the potential cancer driver mechanism for the different enzyme families in general and helicases in particular, we next used WGS-based mutational analysis from the PCAWG consortium, spanning more than 2,600 cancer genomes ^12^. We used the PCAWG-curated list of driver genes^1,12^, and performed an unbiased analysis for each of the enzyme families. We separated the driver mutation by the type of mutation (single nucleotide variants (SNVs), structural variants (SVs) including copy number alterations and whether they were loss-of-function (LoF) or gain-of-function (GoF) mutations). In agreement with the CGC analysis, we found no or very few driver genes in transaminases, lipases and methyltransferases (Fig. 1E). Strikingly, nucleases and helicases were almost exclusively associated with LoF mutations, primarily driven by SVs (grey and dark grey bars). In total, 20 out of 20 (100%) nucleases and 16 out of 17 (94%) helicases were exclusively LoF driver mutations (Fig. 1E). Given the importance of LoF alterations, we next used the larger TCGA dataset to perform a copy number recurrence analysis using GISTIC ^13^, focusing on genes with copy number losses. We found 35 of the 96 (36%) most frequently mutated helicases and nucleases to be associated with a significant GISTIC loss (*P* = 2.8x10^-12^ and *P*=4.15x10^-16^ for helicases and nucleases, respectively, Fisher’s combined probability, Supplementary Table 2), with 31 (89%) of these being significant in two or more cancer types (Fig 1C, red asterisks). In comparison, frequently mutated enzyme families such as proteases, phosphatases and G-protein enzymes were more frequently associated with GISTIC amplification peaks (Supplementary Table 2). Inspired by the potential role of nuclease LoF mutations, we also investigated their alterations at the individual gene level. Indeed, TSGs such as *FANCM* and *DICER* were frequently mutated in a variety of cancers (Extended data Fig. 1B)^14–16^.

Finally, we asked whether specific cancer types were enriched for alterations in helicases and nucleases. Across 18 different cancer types (9,660 patients), we found breast cancer (BRCA) to be highly enriched for helicases, with 151 out of 304 (50%) helicase and nuclease genes to be mutated in at least 2 patients, compared to an average of 54 (18%) helicase and nuclease genes for the other 16 non-skin cancer types (*P*<1x10^-4^, 10,000 permutations, Extended data Fig. 1C). Pathway analysis showed that the helicase and nuclease genes mutated in this cancer subtype were enriched for DNA repair functions, unwinding of DNA, and DNA strand elongation (Extended data Fig. 1D), suggesting deficiencies in genome maintenance.

In summary, we find that helicases and nucleases represent highly mutated enzyme families with the highest proportion of known cancer-driver genes, almost all of which are LoF associated with somatic SVs.

### Genetic screen identifies LoF helicases and nucleases associated with genomic instability

We were motivated by our cancer genomic findings of recurrently mutated helicases and nucleases enriched for DNA repair functions to investigate their potential role in genome maintenance through LoF screening. Therefore, we designed siRNA and CRISPR libraries targeting the 96 most frequently mutated helicase and nuclease genes. We performed image-based phenotypic arrayed screens using siRNA in the osteosarcoma cell line U-2 OS that allows the detection of direct genome instability evident as micronuclei formation ^17,18^.

Further, we carried out siRNA and CRISPR screens in the non-transformed breast epithelial MCF10A iCas9 p53KO cell line^19,20^. Both RNA interference (RNAi) and CRISPR/Cas9 screens were performed in 384-well plate format, targeting each gene individually with three different siRNAs or guide RNAs, respectively. Deficiency in genome maintenance was analysed three days post-transfection by staining for γH2AX and examining micronuclei formation, as established markers of DNA damage ^21,22^ (Fig. 2A-B). In total, we found 9 out of 95 tested genes to be significantly associated with genome maintenance in both the siRNA and the CRISPR screen in the MCF10A iCas9 p53KO model. *Aquarius* (*AQR*) was the top scoring gene followed by the *DEAH-box helicase 8* (*DHX8*) in both the siRNA and CRISPR screens (*P*<0.05, Fisher’s combined p-value, Fig. 2C, Extended Data Fig. 2A-C, Supplementary Table 3). The top-scoring genes *AQR*, *DHX8*, and *DEAH-box helicase 38* (*DHX38*) were validated in independent experiments (Fig. 2D-F and Extended Data Fig. 2D- F). In conclusion, we identified the recurrently mutated helicase genes *AQR*, *DHX8*, and *DHX38* as likely genome maintenance factors. Notably, *AQR* has previously been associated with increased genome instability ^23^ but neither of these three genes were linked with tumour suppression. In line with our findings from the enzyme family-centric analysis, our data showed an enrichment of helicase genes with marked genome maintenance function compared to nuclease genes (Extended Data Fig. 2G).

**Fig. 2.**
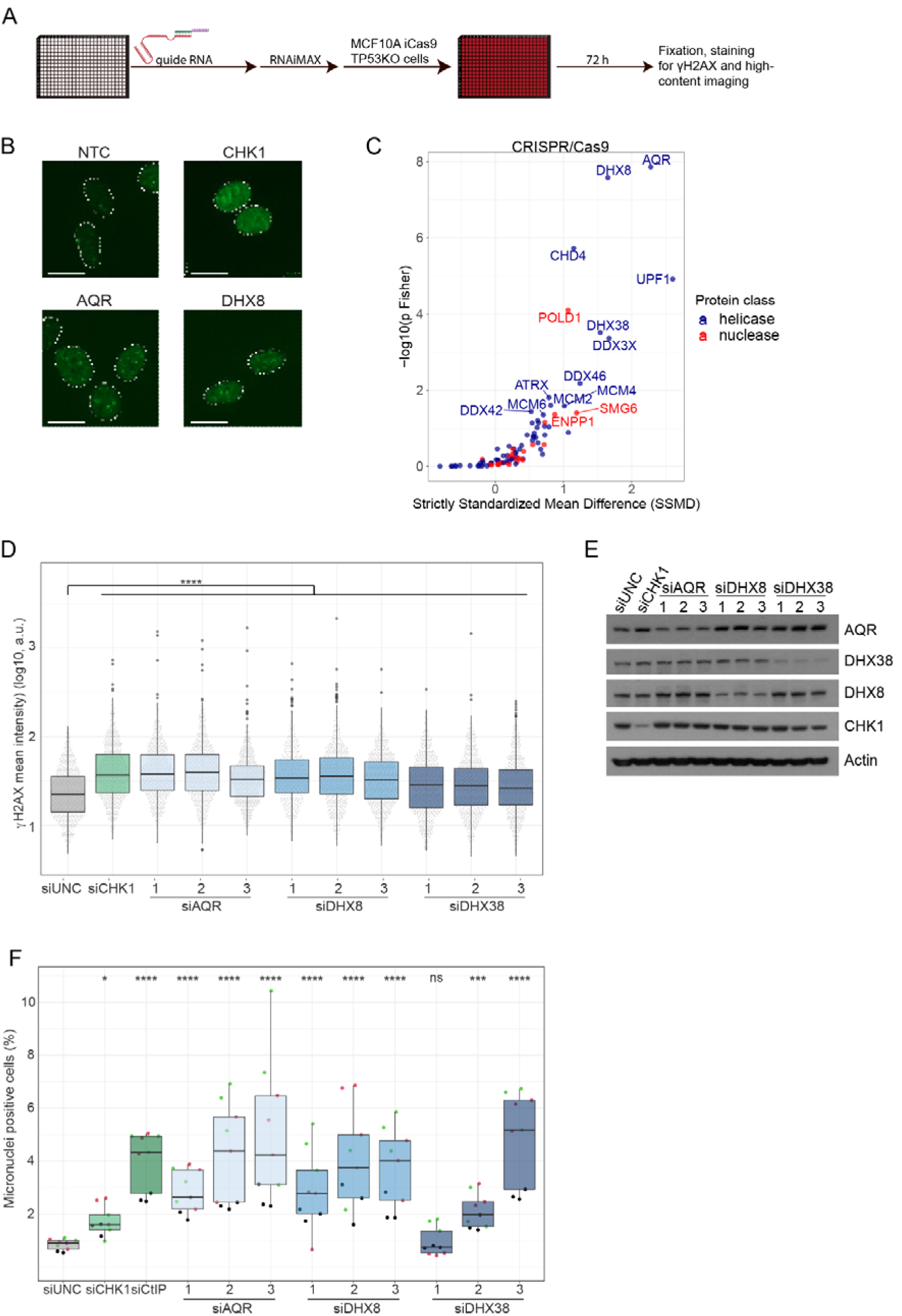
Phenotype screens identifies cancer-associated genome maintenance helicases and nucleases. A) Genome instability screen scheme. Guide RNA complexes were reversely transfected into MCF10A iCas9 p53KO and cells were fixed 3 days post-transfection. Samples were stained for DAPI and γH2AX. B) Representative images of γH2AX, outline indicates outline of DAPI staining, the scale bar represents 20 µm. C) MCF10A iCas9 p53KO CRISPR screen results, performed as shown in A. On the x-axis the effect size is plotted as the strictly standardised mean difference (SSMD) in frequency of γH2AX positive cells. The y-axis corresponds to the Fisher combined p-value from three biological replicates. Coloured by enzyme family and labelled dots represent significantly scoring genes (p Fisher <0.05). D) siRNA screen validation of AQR, DHX8 and DHX38-depleted MCF10A iCas9 p53KO cells. Timing as in A, depicting the γH2AX mean intensity on the y-axis. Representation of one out of three biological replicates. n=600, n=200 per technical replicate. **** *P*<0.0001. E) Immunoblot analysis of siRNA screen validation samples shown in D. Actin was used as a loading control. F) Frequency of micronuclei positive U-2 OS cells of same samples as shown in Extended Data Figure 2D-F. Representation of three biological replicates, each with three technical replicates. Point colours indicate samples belonging to the same biological replicate. n=9.

### AQR hemizygous tumours are associated with homologous recombination deficiency and genome instability

Next, we pursued a pan-cancer analysis of our top-scoring gene *AQR* to further understand its role and influence on cancer genomes and genomic instability. Our initial cancer genomic analysis identified recurrent copy number losses spanning *AQR* (Fig. 1C). In agreement, we found frequent heterozygous copy number loss at *AQR* (Extended Data Fig. 3A), which was associated with increased genomic instability (Extended Data Fig. 3B, P<0.0001) to a level comparable to that of tumours with heterozygous loss of *TP53.* In contrast to *TP53* and *BRCA2*, we found a significant depletion of homozygous *AQR* loss (Extended Data Fig. 3B, P=0.016, Fisher’s exact test), suggesting haploinsufficiency. Motivated by the link between *AQR* loss and copy number genomic instability in breast cancer from our pan-cancer analysis and mammary cell lines (Fig. 1C, Extended Data Fig. 1B-C, Fig. 2), we next used the PCAWG whole-genome sequencing resource to explore whether breast cancer samples with *AQR* loss displayed particular patterns of mutational signatures. We compared breast cancer samples with heterozygous loss of *AQR* (N=38) to cancers with homozygous loss of *BRCA1* (N=7), *BRCA2* (N=15) or *TP53* (N=90). Regressing different mutational signatures on the mutation status of these genes, we found an expected association between *BRCA1* (*P*=6.48x10^-5^, yellow cross) and *BRCA2* (*P*=2.18x10^-5^, red cross) and the mutational signature SBS3 (“BRCAness” signature, Fig. 3A i and Extended Data Fig. 4), which has been found to be elevated in tumours with homologous recombination deficiency (HRD) ^24^. While cancers with heterozygous *AQR* loss or *TP53* homozygous loss alone (*P*=0.012, blue square and *P*=0.005, green cross, respectively) were associated with elevated SBS3, we found the strongest association in samples with both *AQR* and *TP53* loss (*P*=1.75x10^-8^, purple circle, Fig. 3Ai). Our analysis confirmed the previously reported association between *BRCA2* homozygous loss and ID6 (*P*=4.77x10^-12^, Fig.A 3ii and Extended Data Fig. 4) ^25,26^, an indel signature characterized by deletions 5 bp or longer with flanking microhomology. Consistent with our SBS3 signature, we again found *AQR*/*TP53* double mutant cancer samples to have elevated levels of ID6 (*P*=1.92x10^-5^), but even more so for ID8 (*P*=2.49x10^-10^). ID8 is characterized by deletions of five bp or more with no or 1 bp of microhomology and has previously been linked with non-homologous end joining (NHEJ)^26^. We next investigated larger deletions and duplications in size ranges of 0-1kb, 1-10kb, 10-100kb and 100kb-1Mb, which again revealed the expected association between *BRCA2* and short deletion as well as between *BRCA1* and short to medium size duplications (Fig. 3A iii-iv)^27^. While *AQR*/*TP53* double mutant cancers were associated with increased levels of short deletions (1-10kb, *P*=1.51x10^-3^, Fig. 3A iii and Extended Data Fig. 4), the double mutant cancers displayed an even more striking accumulation of short, and medium-sized duplications (1-10kb, *P*=9.70x10^-10^ and 10-100kb, *P*=1.45x10^-7^, Fig. 3A iv and Extended Data Fig. 4)^27^. We also found deletions and duplications in *AQR/TP53* double mutant cancers to have short flanking homologies (0-3 bp, Extended Data Fig. 4). Our analysis identified strong co-occurrence of mutational signatures including ID8, SBS3, ID6, short deletions and duplications in *AQR*/*TP53* double mutant cancers (Fig. 3B, purple bar), but not signatures associated with other mutational processes such as APOBEC (Fig. 3B, SBS13). Importantly, *TP53* homozygous loss cancers without *AQR* heterozygous loss displayed insignificant associations with these signatures of HR and NHEJ (Fig. 3AB, Extended Fig. 3). In support of a tumour-promoting effect, we found significant co-occurrence of *AQR* and *TP53* (*P*< 2.22x10^-16^), while *BRCA1* and *BRCA2* were much less likely to co-occur with *AQR* loss (Extended Data Fig. 5).

**Fig. 3.**
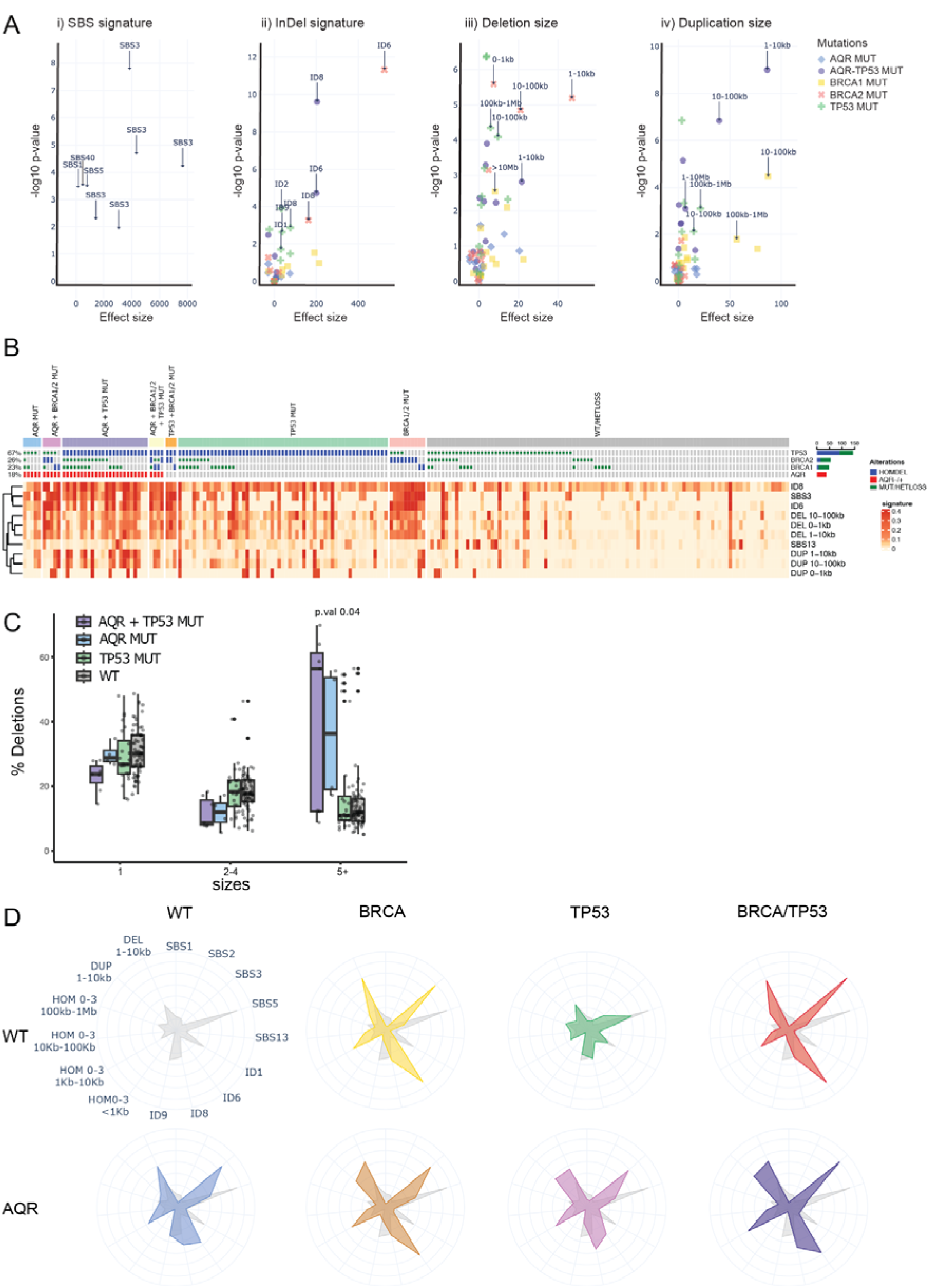
Hemizygous loss of AQR is linked to genome instability and deregulated DNA repair pathways including homologous recombination. A) Statistical testing using a linear model of mutational signatures regressed on the mutational status of hemizygous *AQR* loss, homozygous loss of *TP53,* of *BRCA1* and of *BRCA2* in PCAWG breast cancer samples. The figure shows the enrichment in mutational signature exposure of i) SBS signatures ii) InDel signatures, iii) Deletion sizes and iv) Duplication sizes. Coloured shapes represent the different mutual exclusive mutation types, with e.g. “AQR MUT” representing tumours with heterozygous loss of AQR but no homozygous loss of *TP53*, *BRCA1*or *BRCA2*. B) Oncoprint showing the PCAWG breast cohort stratified by hemizygous *AQR* loss, as well as *TP53*, *BRCA1*, and *BRCA2* homozygous loss. Each column represents a donor, and it displays relative exposures of indels signatures ID8 and ID6, mutational signature SBS3 and SBS13 as well as small duplication and small deletion patterns between 0 to 100Kb. C) Deletions grouped by 1, 2 to 4 or bigger than 5 nucleotide indels in *AQR* heterozygous loss samples (AQR MUT*) AQR* heterozygous loss and TP53 homozygous loss double mutant (AQR+TP53 MUT) and AQR wt and TP53 wt (WT). D) Radar plot showing mutational signature exposure of breast cancer samples with heterozygous *AQR* loss (AQR, bottom row), and comparing to cancers without BRCA or TP53 (WT), with *BRCA1* and/or *BRCA2* homozygous loss (BRCA), *TP53* homozygous loss (TP53) or both BRCA1/2 and TP53 homozygous loss (BRCA/TP53).

Motivated by the strong association with short deletions (ID8), we compared the proportions of deletions of 1bp, 2-4 bp and deletions of length 5 or more (5+, Fig. 3C). We found no difference for cancers with homozygous *TP53* loss alone. Heterozygous *AQR* loss, on the other hand, was associated with a significantly higher proportion of 5+ deletions, which was exacerbated in double mutant *AQR*/*TP53* cancers.

Summarising the different mutational signatures and comparing cancers without *AQR*, *TP53*, *BRCA1*, and *BRCA2* (“WT” in Fig. 3D), we find *AQR* private mutated cancers to show elevated levels of SBS3, ID6, ID8, and 1-10kb deletions (Fig. 3D, blue radar plot), which we find amplified in *AQR*/*TP53* double mutant cancers (Fig. 3D, purple radar). Importantly, *TP53* private mutations does not lead to elevated levels of the investigated signatures compared with WT (Fig. 3D, green radar). *BRCA1* or *BRCA2* mutated cancers (Fig 3D, “BRCA”, yellow radar) display an expected association with especially SBS3 and ID6, which is not notably affected by co-mutations with *AQR* (Fig. 3D. brown radar) or *TP53* (Fig. 3D, red radar).

These findings suggest that homozygous loss of *TP53* potentiates cancers with *AQR* loss to a more extreme somatic phenotype linked with both HRD and NHEJ, characterised by 1- 10kb deletions and duplications, short, 5+ deletions and SBS3.

To further explore the functional role of AQR in genome maintenance and HR, we generated an inducible siRNA-resistant AQR-GFP system in MCF10A cells. We first used γH2AX as a marker for DNA damage to measure the impact of *AQR* depletion in MCF10A cells. We found a significant increase in γH2AX signal upon *AQR* knock-down, which we could rescue using the inducible AQR-GFP system (Extended Data Fig. 6A-B). Next, we assessed HR efficiency upon knocking down AQR using the Direct Repeat GFP (DR-GFP) reporter system cells, which showed almost 2-fold reduced HR efficiency following AQR depletion (Extended Data Fig. 6B-C), in line with a previous report ^28^. Recently, the presence of RNA:DNA hybrids, also called R-loops, was shown to affect HR repair ^29,30^, and AQR has been proposed to be able to resolve R-loops ^31^. For this reason, we tested whether overexpression of RNase H1, an enzyme that degrades the RNA moiety in R-loops, can rescue DNA damage accummulation. Using knock-down of senataxin (SETX) as a positive control, we found that RNase H1 overexpression in AQR-depleted cells reduced levels of the γH2AX marker (Extended Data Fig. 6E-F). This suggests that AQR may maintain genome stability in part through R-loop homoeostasis.

### AQR haploinsufficiency causes S⍰phase-specific damage and sensitivity to PARPi

Our findings suggested a haploinsufficient tumour suppressor role for AQR. To further explore this notion, we generated *AQR* heterozygous MCF10A iCas9 p53KO cells using CRISPR/Cas9 gene editing (Fig. 4A). We were only able to obtain hemizygous but no homozygous knockout clones in agreement with the essential role of AQR in cell survival. *AQR* hemizygosity led to a decrease in AQR protein levels (Extended Data Fig. 4A), and the *AQR* hemizygous cells showed increased γH2AX levels compared to *AQR* diploid cells (Fig. 4B) but no change in cell cycle, as assessed by total DAPI intensity and mean EdU intensity (Extended Data Fig. 7). To further evaluate the impact of *AQR* hemizygosity, we performed low coverage (0.5-1x coverage) whole-genome sequencing of 20 *AQR* hemizygous and 20 *AQR* diploid clones, followed by copy number analysis. We found on average a significant increase in whole-genome instability of the hemizygous *AQR* clones compared to diploid *AQR* clones (*P*=7.8x10^-225^; Fig. 4C). The majority of *AQR* hemizygous clones had three or more large genomic aberrations, compared to approximately 25% of control clones (Fig. 4D). We performed deep whole genome sequencing of a hemizygous *AQR* clone, which demonstrated a locus on chromosome 13 showing highly complex structural variants involving series of small copy number oscillations, reminiscent of the HRD patterns observed in our cancer genomic analysis (Fig. 3). The complex rearrangements at chromosome 13 were connected with additional complex structural variants involving chromosome 6, 7 and X (Fig. 4E-F), suggesting the occurrence of multiple simultaneous DSBs on these chromosomes, followed by illegitimate religation, creating a series of highly connected rearrangements.

**Fig. 4.**
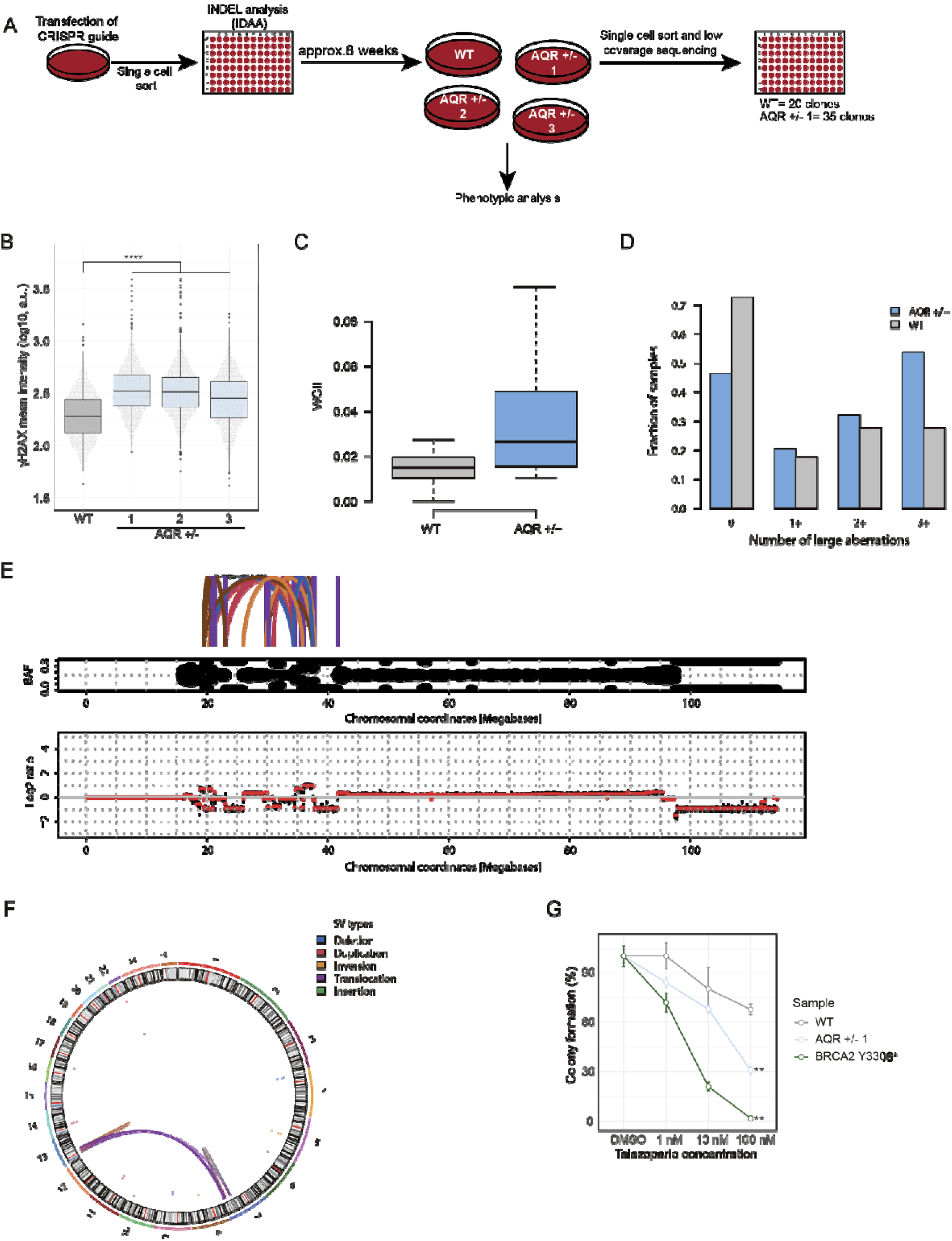
AQR heterozygosity causes increased genome instability and sensitivity to PARP inhibition in a cell line model. A) Schematic presentation of generation and culturing of AQR heterozygous MCF10A iCas9 p53KO cells. B) Analysis of γH2AX mean intensity (y-axis) of WT and AQR heterozygous MCF10A iCas9 p53KO cells. Representation of one out of three biological replicates, n=1500 cells per sample, n=500 per technical replicate. **** *P*<0.0001 compared to AQR WT cells. C) WGII of wildtype clones (grey) and AQR hemizygous depleted clones (blue), p=7.8x10^-225^ D) Paired barplots of wildtype clones (grey) and AQR hemizygous depleted clones (blue) showing the fraction of samples without any copy number alteration (first column from the left), at least 1, 2 or 3 copy number alterations (respectively, second, third and last column from the left) E) Chromosome 13 of an AQR hemizygous depleted clone, showing a catastrophic shattering and rearrangement of the genome. The top of the figure shows arches representing the edges of SV resulting after the rearrangements, the middle of the figure represents B-allele frequency (BAF) binned data, showing loss of heterozygosity when the data is different than 0.5. The bottom panel shows the log2 ratio of the coverage of a wt clone versus the coverage of the mutated clone. positive values represent copy number gains and negative value represents copy number losses F) A circos plot representing the genome-wide scale of structural variants of an AQR hemizygous depleted clone. G) Colony formation of AQR wild type or AQR heterozygous clone 1 MCF10A iCas9 p53KO and BRCA2 Y3308* MCF10A iCas9 cells treated for 2 weeks with the indicated doses of Talazoparib. Representation of technical triplicates within one biological replicate. ** *P*<0.01, compared to the wild-type control.

HR deficiency is typically associated with PARP inhibitor sensitivity. Thus, we sought to test if *AQR* haploinsufficiency sensitises cells to PARP inhibition, potentially posing new treatment options. We employed colony formation assay using *AQR* WT MCF10A iCas9 p53KO, *AQR* hemizygous, and a positive control cell line MCF10A *BRCA2* homozygous Y3308* mutant that is classified as pathogenic by ClinVar (National Center for Biotechnology Information. ClinVar; [VCV000052916.29], https://www.ncbi.nlm.nih.gov/clinvar/variation/VCV000052916.29 (accessed June 3, 2023)). Indeed, the *AQR* hemizygous cells showed sensitivity towards the PARP inhibitor Talazoparib relative to diploid *AQR* cells (Fig. 4G). These results support a role of AQR in HR repair and are in line with the identification of several mutational events associated with HRD in *AQR* hemizygous tumours.

### Helicases have a high prevalence of hemizygous cancer drivers

Our pan-cancer analysis showed that helicases are *i*) frequently mutated, *ii*) almost exclusively TSGs, and *iii*) frequently affected by structural variants and copy number loss. Moreover, our results indicate that *AQR* has a hemizygous cancer driver role with homozygous lethality. In keeping with this, we found *AQR* to be a common essential gene in DepMap^32^, a large-scale functional genomics profiling of cancer dependencies and essential genes. To explore whether recurrent mutations leading to hemizygosity of essential genes represent a broader group of cancer driver genes, we combined mutation recurrence from the pan-cancer analysis of whole genomes (PCAWG) cohort with gene dependencies from the DepMap genome-wide screen. To this end, we found that helicases on average display significantly higher essentiality (Fig. 5A, negative DepMap Dependency score represents the level of essentiality in the CRISPR screen), compared to the other enzyme families.

**Fig. 5.**
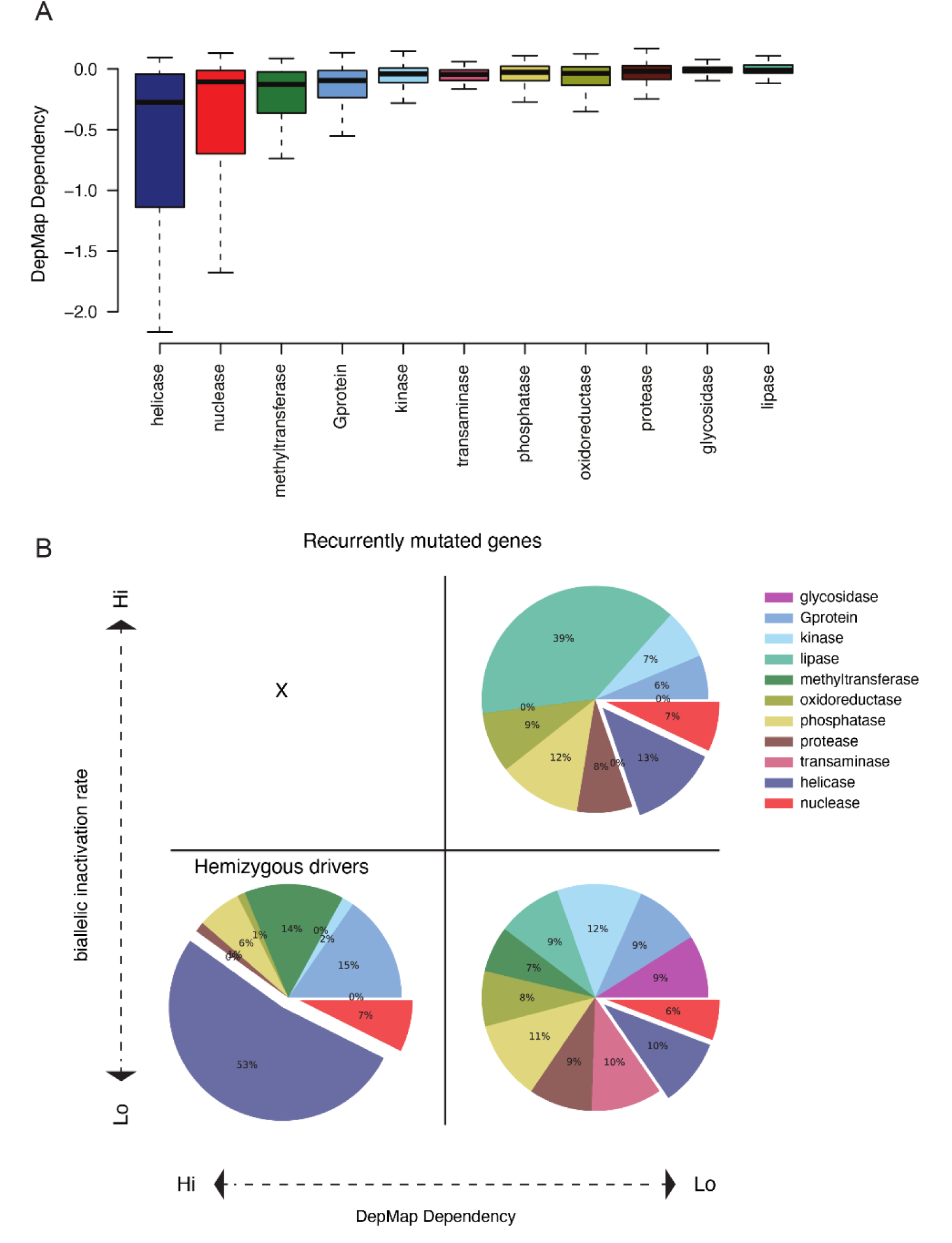
Helicases are enriched for hemizygous driver alterations in DepMap dependence genes. A) Helicase enzymes are highly enriched for dependency genes. DepMap dependency score for the genes in each of the 11 enzyme families based on DepMap CRISPR 22Q4 screen. B) Hemizygous Drivers identified as recurrently mutated genes (at least 3 focal mutations, or a pan-cancer focal recurrence > 0.1%), essentiality (Dependency score < -1) and not found biallelic inactivated (<0.5%).

We next used a simple schema to classify hemizygous driver candidate genes, requiring gene dependence essentiality (DepMap dependency score < -1), recurrently mutated (requiring at least three focal somatic alterations and a focal mutations rate > 0.1% in the cohort) and strongly depleted for biallelic inactivation (biallelic rate < 0.1%). We separated genes and their associated enzyme family into four groups, *i*) non-essential, hemizygous (Fig. 5B, lower right), *ii*) non-essential, biallelic inactivation,(representing well-known TSGs such *TP53* and *PTEN,* Fig. 5B, top, right), *iii*) essential and biallelic inactivated, which as expected never occurred (Fig. 5B, top left), and *iv*) essential, hemizygous (Fig. 5B, lower left). We found in total 42 putative hemizygous cancer drivers, 23 of which were helicases including *AQR* (55%, *P*=1x10^-12^, Fisher’s exact test, Supplementary Table 4). Helicases were underrepresented among non-recurrently mutated genes (Extended Fig. 8A). In support of critical roles in nucleic acid maintenance and processing, the helicase hemizygous driver candidates were enriched for genes involved in DNA and RNA processes such as DNA replication, Spliceosome and NHEJ (Extended Fig. 8B).

In summary, we identify a subset of putative hemizygously mutated TSGs highly enriched for helicases. This group is characterised by frequent heterozygous losses but never biallelic inactivation, corroborated by high essentiality, suggesting important cellular functions.

## Discussion

Though proteins have multiple roles in cellular and organismal processes, cancer driver functions are frequently linked with their catalytic activities. We based our study on this notion and with a focus on enzyme function, we uncovered substantial differences in cancer mutation rate across the eleven different enzyme families. While prior focus has been on elucidating especially kinases as drivers in cancer, we discovered a surprisingly high mutation rate in helicases, a family of enzymes involved in processes including transcription, replication, translation and DNA repair. We demonstrate that helicases are a predominant family of cancer drivers and that somatic alterations affecting helicases are almost exclusively associated with LoF. In agreement, we found helicases to be frequently disrupted by SVs including copy number losses, leading to hemizygosity, which we suggest is predicated on haploinsufficiency and the common essential roles of a subset of helicases.

Knudsen’s classical two-hit hypothesis has been used to guide the discovery of TSGs in cancer, but a large proportion of the cancer genome is undergoing recurrent hemizygous loss. Several TSGs have been associated with a haploinsufficient role in cancer including *SMAD4*, *TP53*, *CDKN1B*, *NF1* and *PTEN* ^33–35^. Notably, these haploinsufficient genes all display frequent biallelic inactivation, associated with a more severe cancer-driving mechanism ^12,36,37^. Here, we extend this concept and provide evidence for the presence of a subset of hemizygous essential TSGs particularly frequent in helicases. Given the important role of helicases in essential processes involving the unwinding of nucleic acids, it is tempting to speculate that the helicase protein levels are finely tuned to operate in these vital processes and therefore sub-lethal but vulnerable to reduced dosage upon hemizygous loss, leading to cell survival but with genomic instability under hemizygous conditions.

The nuclease family was also enriched for LoF cancer drivers, though mutation frequency was lower than for helicases. Both helicases and nucleases play important roles in DNA repair and suppression of genome instability in cancer ^38,39^ and we show that they are almost exclusively undergoing genomic losses rather than gains in cancer genomes. This motivated us to pursue functional LoF screening of the most frequently mutated helicases and nucleases from our pan-cancer analysis, and it enabled us to extend, compare and functionally test the cancer-promoting or suppressing function of multiple potential driver genes ^40–43^.

Among the 96 candidate helicases and nucleases, we noted that AQR suppression displayed a particularly strong phenotype associated with genome maintenance. Moreover and in keeping with a hemizygous role, we leverage a large pan-cancer whole genome sequencing dataset to show three striking features of AQR. First, *AQR* is exclusively hemizygously deleted and never biallelic inactivated. Second, cancer genomes with hemizygous *AQR* loss frequently co-occur with homozygous *TP53* loss, and we provide support that *TP53* potentiates the genomic consequences of AQR loss, including genomic signatures compatible with NHEJ and HRD in breast cancer genomes. Third, clonogenic experiments with hemizygous loss of *AQR* result in similar complex somatic aberrations and it is associated with sensitivity towards PARP inhibitors. We note that *RAD51*, a gene involved in HRD and found to interact with BRCA1 and BRCA2 ^44,45^, is located approximately 6 Mb downstream of *AQR*, and is frequently lost in conjunction with AQR losses. Loss of both *AQR* and *RAD51* may be additive in contributing to the observed HRD-related mutational signatures, but we find compelling support for an AQR-private role. First, our CRISPR and siRNA-based screens specifically target *AQR*, and loss of *AQR* was associated with a strong phenotype associated with genomic instability. Second, cloning of hemizygous CRISPR-engineered *AQR* cells led to an accumulation of somatic alterations including complex SVs. Finally, one of the breast cancer samples with a strong signature exposure of both SBS3, ID6, ID8 and small deletions had a small genomic deletion that encompassed *AQR* but not *RAD51* (Extended Figure 4B, second from right in the “AQR MUT” group), supporting an *AQR*-specific phenotype.

In the *AQR* hemizygous tumour samples, genome instability and HR-deficiency related signatures might be caused by multiple mechanisms. AQR has been linked with RNA splicing and R-loop suppression^31,46^, and AQR was suggested to promote HR in response to different double-strand break-inducing agents ^28^. Collectively, these functional perturbations may jointly underlie genome instability in *AQR* hemizygous cells. *AQR* is a common essential gene, and targeting such essential genes therapeutically can prove difficult due to high toxicity, though it is notable that the drugging of essential kinases such as WEE1, ATR, and CHK1 are being explored in multiple clinical trials ^47,4847^. Thus, it is interesting to speculate that cells with hemizygous loss of *AQR* could be more vulnerable to AQR-targeted therapies than normal diploid cells. More directly, our findings also support a beneficial effect of PARP inhibitor treatment of *AQR* haploinsufficient tumours. Altogether, we propose that subset of helicases are driving cancer development through a mechanism involving hemizygous loss, and that the affected pathways present targetable cancer vulnerabilities.

Functional-mechanistic experiments and larger cancer-focused analyses will be needed to explore and expand the role and implication of hemizygous drivers in cancer development and treatment.

## Acknowledgements

We thank Elin Pietras, Cornelia Steinhauer and Henning Gram Hansen from the high-throughput cell-based screening facility at BRIC for their assistance. We thank Zoltan Szallasi and members of the JW and CSS labs for manuscript comments.

CSS is funded by the Danish Cancer Society, Danish Medical Research Council, and Lundbeck Foundation. CSS and KVO have received funding from the European Union’s Horizon 2020 research and innovation program under the Marie Sklodowska-Curie grant (agreement number 722729). JW is funded by the Danish Cancer Society, Novo Nordisk Fonden and Danish Medical Research Council. FF is funded by Novo Nordisk Fonden. BRM was funded by Merck KGaA, Darmstadt, Germany (CrossRef Funder ID: 10.13039/100009945). BRM present address is Research Unit Oncology, Merck KGaA, Darmstadt, Germany.

## Materials and Methods

### Cell culture

The human osteosarcoma cell line, U2-OS, was cultured in Dulbecco’s modified Eagle Medium (DMEM) supplemented with 10 % FBS (HyClone, HYCLSV30160.03) and 1 % penicillin/streptomycin (GIBCO, 15140-130). The human breast epithelial cell line MCF10A was cultured in DMEM F-12 (GIBCO, 31330095) supplemented with 5 % horse serum (GIBCO. 26050088), 1 % penicillin/streptomycin (GIBCO, ), 10 μg/ml Insulin (Sigma Aldrich, I1882-100MG), 0.5 μg/ml Hydrocortisone (Sigma Aldrich, H0888-1G), 20 ng/ml EGF (PeproTech, AF-100-15) and 100 ng/ml Cholera toxin (Sigma/Merck, C8052-5MG).

### CRISPR and siRNA screens

Cherry□pick libraries targeting the 96-selected helicase and nuclease genes were purchased from Sigma (siRNA) and Dharmacon (crRNA) containing three siRNAs or guide RNAs to target each gene, respectively. Location in the 384-well plate of crRNAs was randomised using an ECHO liquid dispenser to avoid location bias. The screens were performed using a Hamilton STARlet liquid dispenser. For the CRISPR screen, Cas9 expression was induced 24 h prior to transfection using 1 μg/ml Doxycycline (Dox). MCF10A iCas9 p53KO or U□2 OS cells were reversely transfected with 10 nM silencer select siRNA or 25 nM crRNA:tracrRNA complex. 72 h post□transfection, cells were fixed in 4 % Formaldehyde (VWR) for 15 min, followed by four washes with PBS. Afterwards, cells were permeabilized in 0.25 % Triton□X 100 for 10 min, washed four times with PBS before blocking in 3 % BSA for one hour at room temperature. Incubation with the primary antibody targeting γH2AX (Merck Millipore 05□636, 1:1000) was performed overnight. The following day, four PBS washes were performed followed by 1 h incubation at room temperature with the anti-mouse Alexa Fluor 488 antibody (Invitrogen, A-21202), four PBS washes, 30 min incubation with 1 μg/ml DAPI and five final washes with PBS. 16 imaging fields were acquired with an IN Cell Analyzer 2200 microscope and a 20x objective (GE Healthcare) to obtain ∼2000 cells/well for control samples for the analysis with the IN Cell Analyzer Workstation (Cytiva) software. Nuclei were segmented based on DAPI staining using the tophat segmentation method and the mean γH2AX intensity was measured within the nuclei.

The data analysis was performed as described in^49^, based on the percentage of γH2AX positive cells per well and biological triplicates. In addition, the Strictly Standardised Mean Difference (SSMD) was calculated per gene, based on the normalised levels of γH2AX positive cells to the negative control^50^.

### siRNA transfections

Cells were seeded the day before transfection with 50 nM siRNA using Lipofectamine RNAiMAX (Thermo Fisher Scientific, 13778500). The media was changed or cells were reseeded 5 h post□transfections. The MISSION siRNA Universal Negative Control #1 (siUNC, Sigma Aldrich, SIC001-10NMOL) was used as a negative control. All siRNA sequences can be found in Supplementary Table 3.

Generation of p53KO, heterozygous AQR and BRCA2 3308 mutation cell lines MCF10A iCas9 cells were transfected with an all-in-one pSpCas9(BB)-2A-GFP plasmid (PX458, Addgene 48138), containing a CRISPR guide sequence targeting TP53 exon 2 (TCGACGCTAGGATCTGACTG). Cells were sorted for medium Cas9-GFP expression using a FACS Aria III (BD) and treated with 10 μM Nutlin3a (Tocris Bioscience, 6075) to enrich for clones with p53 knockout. Single-cell clones were screened for p53 knockout by Indel Detection Amplicon Analysis (IDAA) and validated by sequencing^51^. Twelve clones, each with a homozygous loss of p53, were pooled and used for further experiments.

To generate MCF10A iCas9 p53KO cells heterozygous for AQR, cells were seeded and Cas9 expression was induced by treatment with 1 μg/ml Doxycycline. The following day, cells were transfected with 15 nM crRNA (UAGGCCUUGCUAGAUACCUC):tracrRNA complex using Lipofectamine RNAiMAX. Two days post-transfection, cells were single□cell sorted into 96□well plates using a FACS Aria III (BD). Indel formation was analyzed by IDAA and cells with a frameshift indel as well as a wild□type clone as an internal control were selected for further testing. The initial passage (p1) was used for phenotypic evaluation, sequencing and was cultured for 30 passages (p31) over 15 weeks and treated for 16 h with 0.5 µg/ml Aphidicolin once a week.

To generate BRCA2 Y3308* mutation in MCF10A iCas9 cells, cells were seeded and Cas9 expression was induced by treatment with 1 μg/ml Doxycycline 24 hours in advance. The following day, cells were transfected with 20 nM crRNA (UAGGCCUUGCUAGAUACCUC):tracrRNA complex using Lipofectamine RNAiMAX together with 10 nM single strand nucleotide donor carrying the mutation. Two days post-transfection, cells were single-cell sorted into 96□well plates using a FACS Aria III (BD). The knock□in formation was analyzed by restriction enzyme digestion, with a single site introduced by the donor, as well as a wild-type clone as internal control was selected for further testing.

Library preparation for whole genome sequencing AQR wild type and heterozygous clones from p1 and p31 were single-cell sorted into 96-well plates using a BD flow cytometer. Single-cell clones were grown up and cell pellets with 500,000 cells were frozen for sequencing. High molecular weight DNA was extracted using Gentra Purigene Cell kit (Qiagen, 158767) following manufacturer instructions for culture cells. Briefly, cells were lysate using 300 µl of Cell Lysis Solution, followed by RNase treatment. After 5 minutes incubation at 37°C, 100 µl Protein Precipitation Solution was added to lysate and remove proteins. After centrifuge for 1 minute at 15,000 g, precipitated proteins form a tight pellet. The supernatant was transferred into a clean 1.5 ml microcentrifuge tube with 300 µl isopropanol. After mixing and spin down, DNA was visible as a white pellet. Final wash with 300 µl of 70% ethanol was done to remove remain impurities. DNA was resuspended in 100 µl DNA Hydration Solution and incubated at 65°C for 1 h to dissolve it. DNA concentration was measured by Qubit dsDNA BR Assay Kit (Thermo Fisher Scientific, Q32850), purity by Denovix spectrophotometer (DS-11Fx) and DNA integrity by electrophoresis in 0.8% agarose gel.

We performed shallow WGS on 113 cell line clones, using 400 ng of genomic DNA to prepare PCR free libraries using the Illumina DNA PCR-free kit (20041794). This strategy provides several benefits for accurate sequencing i.e.: eliminate PCR duplicates and polymerase errors. It combines tagmentation on beads with extension using IDT for Illumina DNA Unique Dual Indexes (Illumina DNA PCR-Free Library Prep, Reference Guide 1000000086922 v01). Final libraries were quantified by Qubit ssDNA Assay Kit (Thermo Fisher Scientific, Q10212), and size distribution was assessed by Bioanalyzer High Sensitivity DNA Kit (Agilent, 5067-4626). Libraries were combined at 2nM concentration pool and pair end sequenced using NextSeq 500/550 High Output Kit v2.5, 150 cycles, (Illumina, 20024907).

As a validation, deep whole genome sequencing was done on six randomly selected samples. Libraries were prepared using 200 ng of genomic DNA following the NEBNext Ultra II FS DNA Library Prep Kit for Illumina (New England Biolabs, E7805S). Briefly, DNA was enzymatically fragmented for 12 minutes at 37°C to get 450 bp fragments on average. After size selection, end repair, dA-tailing and UDI Adaptor Ligation, final libraries were amply by 5 PCR cycles, and quantified by Qubit dsDNA High Sensitivity assay Kit (Thermo Fisher Scientific, Q10212). Size distribution was assessed by Bioanalyzer High Sensitivity DNA Kit (Agilent, 5067-4626). Finally, libraries were combined at 2nM concentration pool and pair end sequenced using 300 cycles on NovaSeq 6000 S2 Reagent Kit v1.5, 300 cycles, (Illumina, 20028314). Variant calling was performed as described in ^52^.

### Generation of AQR-GFP inducible MCF10A cells

For lentivirus production, HEK293T cells were transfected with 3 μg pLVX, 1 μg VSV-G (Clontech) and 1 μg PAX2 (Clontech) plasmids using JetPEI (Polyplus, 101-10N) according to the manufacturers’ protocol. 5 h post-transfection the transfection media was aspirated and cells were cultured in 10 ml MCF10A media. Two days after plasmid transfection, lentiviral particles were collected by centrifugation of the supernatant at 300 g for 5 min. For the transduction, 7 ml of supernatant was mixed with 3 ml MCF10A culture media, and polybrene (Sigma Aldrich H9268-5G) was added at a final concentration of 10 μg/ml.

MCF10A cells were transduced on two consecutive days and cells were selected with 3 μg/ml Puromycin. In addition, AQR-GFP expression was induced with 1 μg/ml Doxycycline for 48 h, and cells were sorted for GFP expression using a FACS Aria III (BD). Different GFP expression levels were tested by WB to match with endogenous AQR expression. The pLVX plasmid containing only the GFP sequence was used to generate GFP inducible expressing cells.

### R□loop testing by RNase H1 overexpression and DRB treatment

U□2 OS cells were transfected with siRNAs as described above followed by reseeding and 48 h later plasmid transfection of either pEGFP or pEGFP-RNase H1 (M27) plasmids^53^ using GenJet (SignaGen Laboratories, SL100489-OS) according to the manufacturers’ protocol. 24 h post plasmid transfection, cells were harvested and prepared for immunofluorescence or immunoblotting as described below. γH2AX mean intensity was measured for GFP-positive cells only.

To assess the effect of transcription inhibition on genome instability, MCF10A iCas9 p53KO cells were transfected with siRNA and after 66 h treated with 150 μM DRB (5,6-dichloro-1- beta-D-ribofuranosylben- zimidazole, Sigma, D1916-10MG) for 6 h. Samples were prepared for immunofluorescence and immunoblotting.

### Homologous recombination assay

U□2 OS cells transfected with pDR-GFP plasmid (Addgene, 26475) using GenJet (SignaGen Laboratories, SL100489-OS) according to the manufacturer’s protocol were selected with Puromycin (5 μg/ml). To investigate HR efficiency, cells were seeded in a 6□well plate and the next day transfected with siRNA followed by a media change 5 h post□transfection. 24 h post siRNA transfection, cells were transfected with I□Sce1 plasmid using GenJet according to the manufacturer’s protocol. Two days later, cells were trypsinized and fixed in 1 % formaldehyde for 15 min, followed by PBS wash and permeabilization in 60 % EtOH for 1 h on ice. Afterwards, cells were washed and stained with PI in the presence of RNase A for 30 min at 37 degrees. Samples were analyzed on a FACSCalibur using the Cellquest Pro (BD Biosciences) and FlowJo software. The HR efficiency was calculated for 20,000 cells per sample based on the percentage of GFP-positive cells in S- and G2□phases of the cell cycle, and normalized to siUNC transfected cells.

### PARP inhibitor sensitivity assay

For each sample, 1000 cells were seeded in a well of a 6□well plate, using the AQR wild type or heterozygous clones and a BRCA2 3308* pathogenic variant in MCF10A iCas9 cells as a positive control. Three technical replicates were performed to account for seeding differences. Two days after seeding cells, cells were treated with the indicated doses of Talazoparib or DMSO for 4 days. Afterwards, pre-treated cells were counted and reseeded at a density of 200 cells/well. After another two days, cells were treated again with the indicated doses of Talazoparib for another 6 days. Cells were fixed in 100 % methanol for 10 min, followed by staining with MeOH/Crystal Violet for another 10 min. Colonies were manually counted and numbers were normalized to DMSO-treated cells.

### Immunoblotting

RIPA buffer (Sigma Aldrich, R0278) supplemented with 1% vol/vol aprotinin, 5 μg/ml leupeptin, 1x cOmplete EDTA-free protease inhibitor cocktail (Sigma, 5056489001), 5 mM NaF, 20 mM beta-glycerophosphate and 0.2 mM Na_3_VO_4_ was used to lyse cells on ice. A 102C CE Converter, Branson, was used to sonicate all samples followed by centrifugation at 20,000 g for 10 min. The protein concentration was measured by Bradford assay. SDS-PAGE was used to resolve proteins and proteins were transferred to a nitrocellulose membrane. Membranes were blocked in 5 % milk in PBST for 1 h at room temperature followed by incubation with the primary antibody overnight. Prior to incubation with anti-rabbit or anti-mouse secondary HRP-conjugated antibodies (Vector Laboratories, PI□1000 and PI□2000), and afterwards membranes were washed for 30 min in PBS□T. Information about antibody manufacturer, catalogue number and used dilution can be found in Supplementary Table 3.

### Immunofluorescence

If applicable, cells were pulsed with 10 μM EdU (Thermo Fisher Scientific, A10044) for 30 min before fixation with 4 % formaldehyde for 12 min. After three PBS washes, cells were permeabilized in 0.25 % Triton□X 100 in PBS for 10 min and again washed three times with PBS. Cells were blocked in 3 % BSA in DMEM containing 10 % FBS for 20 min at room temperature. To visualise EdU incorporation the Click-IT cell reaction buffer kit was used according to manufacturer instructions (Thermo Fisher Scientific, C10269) together with Alexa Fluor 647 Azide (Thermo Fisher, A10277). Primary and secondary antibodies were prepared in blocking solution and incubated for 1 h at room temperature. To stain nuclei, DAPI (Sigma, D9542-5MG) was incubated together with the secondary antibody at a final concentration of 0.1 μg/ml. Imaging was performed using an Olympus ScanR workstation and the respective software was used for the analysis.

### immunofluorescence Statistics

All immunofluorescence-based analyses of γH2AX mean intensity were performed in three biological replicates, each containing technical triplicates. All graphs represent one biological replicate due to a large amount of data. Each technical replicate contributes equally to the panels represented in the figures. E.g. n=450 means that 150 nuclei were randomly selected from one technical replicate.Non-parametric statistical analyses were performed on a rank that was assigned to each nuclei based on its γH2AX mean intensity. A linear mixed model was fitted for the ranks, and the technical replicates were used as a variance component to account for biological heterogeneity within an experiment. Multiple comparisons were performed using the lsmeans and the contrast function of the lsmeans package in R (https://www.rdocumentation.org/packages/lsmeans/versions/2.27-2/topics/lsmeans). *P* values were adjusted using the Holm method. Micronuclei analysis was performed as described above but included data from biological as well as technical replicates. The DR-GFP assay was analyzed using a Mann-Whitney U test to compare siUNC to either siAQR or siBRCA2 transfected cells. PARP inhibitor sensitivity was analyzed using a two-way ANOVA test for repeated measures and Dunnet’s multiple comparisons. *P* values are reported with the following significance codes: \*\*\*\**P* < 0.0001, \*\*\**P* < 0.001, \*\**P* < 0.001, \**P* < 0.05.

### Site directed mutagenesis

siRNA resistant AQR plasmids for lentivirus production were generated using site□directed mutagenesis followed by InFusion. As a template, the AQR-GFP plasmid (RG220742) was used for the mutagenesis with KOD polymerase (Merck/Millipore, 71086□3), used according to the manufacturer’s protocol. The siRNA□resistant AQR sequence was cloned into the pLVX-TetOne vector (Clontech Laboratories), containing an N-terminal GFP, using the InFusion HD cloning kit (Clontech Laboratories, 639691) according to the manufacturer’s protocol. Primer sequences can be found in Supplementary Table 3.

### Somatic variant analysis of pan-cancer data

PCAWG data analysis is based on 2,581 WGS profiled tumour samples from the PCAWG consortium ^12^ with 31 histology types (pan-cancer) and 212 unique breast cancer patient samples. Analysis of genomic aberrations was conducted using the official release set of somatic point mutation, copy number alterations and structural variants (SVs) ^54^. Every gene was annotated as mutant if one or several of the following requirements were satisfied, i) nonsynonymous somatic or germline SNV in the gene, ii) somatic copy number loss overlapping at least 50% of the gene body, and/or iii) SV predicted to disrupt the open reading frame of the gene in question. A series of analyses was performed for SV breakpoint recurrence. SV breakpoint was annotated with length of nearby sequence homology, binned by 0-3 nt, 4-11 nt and 12-1000 nt. We computed the proportion of each group, by normalizing to the total number of SVs in each sample. Analysis of deletion and tandem duplication (TD) size distributions was performed, by counting the number of deletions and the number duplications in binned sizes of 1-1Kb, 1Kb-10Kb, 10Kb-100Kb, 100Kb-1000Kb and >1Mb and normalized to the total number of TDs. SBS signature exposure was computed by normalizing the number of inferred SNVs attributed to that SBS to the total number of SNVs for each tumour sample.

Genome instability from TCGA and PCAWG were computed as the percentage of the aberrant genome using the weighted genome instability index (WGII) in which the portion of the aberrant genome is calculated by chromosome, and the mean proportion of the aberrant genome is calculated across the chromosomes. This approach prevents the inflation of the score caused by aberrations in larger chromosomes.

We used the PCAWG segmentation release data including ploidy and purity estimates. We defined the aberrant segments when the copy number estimated for the segment was different from the estimated ploidy, and used the cumulative aberrant segments size to estimate the proportion of aberration relative to the chromosome size.

For TCGA data we used publicly available segmented data from SNP arrays (cBioPortal), we computed the genome mean logR value from the segment data and estimated aberrant segments those outside the logR range of -0.2 and 0.2. We then calculated the WGII score in a similar manner than the PCAWG cohort.

### Mutation enrichment per enzyme family

We collected publicly available mutation data from TCGA (https://portal.gdc.cancer.gov June 8th 2017), comprising Bladder carcinoma (BLCA 99 patients), Breast carcinoma (BRCA 892 patients), Colorectal adenocarcinoma (COAD 233 patients), Esophageal adenocarcinoma (ESCA 141 patients), Head and neck carcinoma (HNSC 384 patients), Kidney clear cell carcinoma (KIRC 417 patients), Kidney renal papillary cell carcinoma (KIRP 291 patients), Low-grade glioma (LGG 515 patients), Liver Hepatocellular Carcinoma (LIHC 377 patients), Lung adenocarcinoma (LUAD 405 patients), Lung squamous cell carcinoma (LUSC 178 patients), Ovarian carcinoma (OV 316 patients), Pancreatic adenocarcinoma (PAAD 185 patients), Prostate adenocarcinoma (PRAD 499 patients), Rectum adenocarcinoma (READ 167 patients), Melanoma (SKCM 118 patients),Thyroid Cancer (THCA 503 patients), Thymoma (THYM 124 patients).

We identified the genes belonging to the seven enzyme classes (namely oxidoreductases, transferases, hydrolases, lyases, isomerases, ligases and translocases) from UniProt and 12 enzyme families (namely glycosidase, Gprotein, kinase, lipase, methyltransferase, oxidoreductase, oxygenase, phosphatase, protease, transaminase, helicase and nuclease) as classified in pantherdb (http://www.pantherdb.org). 66 out of 69 oxygenases genes were redundant with oxidoreductases, and we therefore excluded this family from further analyses due to the low number of unique genes. We computed the number of missense and nonsense mutation, splice site and frameshift deletions for each gene using the TCGA cohort. As an additional criterion, we considered only genes that were mutated in at least two different patients in the same cancer type.

We used a permutation approach to compute the probability of observing a given number of mutated genes within a similar size group of genes randomly selected, repeating the process 10,000 times, for each enzyme class, family and for each cancer type. We calculated the empirical p-value given by the time an equal or higher number of genes were found mutated with respect to the observed number of mutations, divided by the number of iterations.

For each cancer type enzyme class/family combination, we also computed the standardized mean difference, subtracting the mean of the mutated genes in the permutations to the number of observed mutated genes, and dividing by the standard deviation of the permutations.

We computed a single p-value for each enzyme class/family using the Fisher combination test on the empirical p-value for each cancer type.

To obrain an aggregate effect size estimate for each enzyme class/family we calculated the mean of the standardized mean difference for all cancer type. We utilized the information of the 5 and 95 percentiles, as a measure for cancer-type specific effect sizes.

### Permutation-based significance assessment of cancer mutation data

In order to have a background model to compare various analyses on the PCAWG cohort, we applied the curveball algorithm^55^ to the binary matrix of the mutational data, in which columns represent genes and the rows represent patients.

The curveball algorithm constrains the randomization to scenarios where the cumulative sums of each column and row are always constant, providing a background model closer to a real scenario compared to an unbounded or partially bounded shuffling methodology.

We applied the method to obtain 10,000 permuted versions of the mutation binary matrix.

## References

1. Rheinbay, E. et al. Analyses of non-coding somatic drivers in 2,658 cancer whole genomes. Nature 578, 102–111 (2020).

2. Martincorena, I. et al. Universal patterns of selection in cancer and somatic tissues. Cell 171, 1029–1041.e21 (2017).

3. Reyna, M. A. et al. Pathway and network analysis of more than 2500 whole cancer genomes. Nat. Commun. 11, 729 (2020).

4. Leiserson, M. D. M. et al. Pan-cancer network analysis identifies combinations of rare somatic mutations across pathways and protein complexes. Nat. Genet. 47, 106–114 (2015).

5. Schomburg, I. et al. BRENDA in 2013: integrated reactions, kinetic data, enzyme function data, improved disease classification: new options and contents in BRENDA. Nucleic Acids Res. 41, D764–72 (2013).

6. McDonald, A. G. & Tipton, K. F. Enzyme nomenclature and classification: the state of the art. FEBS J. 290, 2214–2231 (2023).

7. Stebbing, J. et al. The regulatory roles of phosphatases in cancer. Oncogene 33, 939– 953 (2014).

8. Bhullar, K. S. et al. Kinase-targeted cancer therapies: progress, challenges and future directions. Mol. Cancer 17, 48 (2018).

9. Morgat, A. et al. Updates in Rhea - an expert curated resource of biochemical reactions. Nucleic Acids Res. 45, D415–D418 (2017).

10. Curtis, C. et al. The genomic and transcriptomic architecture of 2,000 breast tumours reveals novel subgroups. Nature vol. 486 346–352 Preprint at (2012).

11. Thomas, P. D. et al. PANTHER: a library of protein families and subfamilies indexed by function. Genome Res. 13, 2129–2141 (2003).

12. 12. The ICGC/TCGA Pan-Cancer Analysis of Whole Genomes Consortium. Pan-cancer analysis of whole genomes. Nature 578, 82–93 (2020).

13. Mermel, C. H. et al. GISTIC2.0 facilitates sensitive and confident localization of the targets of focal somatic copy-number alteration in human cancers. Genome Biol vol. 12 R41 Preprint at (2011).

14. Zhang, B. et al. A dosage-dependent pleiotropic role of Dicer in prostate cancer growth and metastasis. Oncogene 33, 3099–3108 (2014).

15. Lopes, J. L., Chaudhry, S., Lopes, G. S., Levin, N. K. & Tainsky, M. A. FANCM, RAD1, CHEK1 and TP53I3 act as BRCA-like tumor suppressors and are mutated in hereditary ovarian cancer. Cancer Genet. 235–236, 57–64 (2019).

16. Kiiski, J. I. et al. Exome sequencing identifies FANCM as a susceptibility gene for triple-negative breast cancer. Proc. Natl. Acad. Sci. U. S. A. 111, 15172–15177 (2014).

17. Mackenzie, K. J. et al. cGAS surveillance of micronuclei links genome instability to innate immunity. Nature 548, 461–465 (2017).

18. Hatch, E. M., Fischer, A. H., Deerinck, T. J. & Hetzer, M. W. Catastrophic nuclear envelope collapse in cancer cell micronuclei. Cell 154, 47–60 (2013).

19. Tait, L., Soule, H. D. & Russo, J. Ultrastructural and immunocytochemical characterization of an immortalized human breast epithelial cell line, MCF-10. Cancer Res. 50, 6087–6094 (1990).

20. Soule, H. D. et al. Isolation and characterization of a spontaneously immortalized human breast epithelial cell line, MCF-10. Cancer Res. 50, 6075–6086 (1990).

21. Fernandez-Capetillo, O., Lee, A., Nussenzweig, M. & Nussenzweig, A. H2AX: the histone guardian of the genome. DNA Repair 3, 959–967 (2004).

22. Terradas, M., Martín, M. & Genescà, A. Impaired nuclear functions in micronuclei results in genome instability and chromothripsis. Arch. Toxicol. 90, 2657–2667 (2016).

23. Paulsen, R. D. et al. A genome-wide siRNA screen reveals diverse cellular processes and pathways that mediate genome stability. Mol. Cell 35, 228–239 (2009).

24. Davies, H. et al. HRDetect is a predictor of BRCA1 and BRCA2 deficiency based on mutational signatures. Nat. Med. 23, 517–525 (2017).

25. Thatikonda, V. et al. Comprehensive analysis of mutational signatures reveals distinct patterns and molecular processes across 27 pediatric cancers. Nat Cancer 4, 276–289 (2023).

26. Alexandrov, L. B. et al. The repertoire of mutational signatures in human cancer. Nature 578, 94–101 (2020).

27. Nik-Zainal, S. et al. Landscape of somatic mutations in 560 breast cancer whole-genome sequences. Nature 534, 47–54 (2016).

28. Sakasai, R. et al. Aquarius is required for proper CtIP expression and homologous recombination repair. Sci. Rep. 7, 13808 (2017).

29. Sessa, G. et al. BRCA2 promotes DNA-RNA hybrid resolution by DDX5 helicase at DNA breaks to facilitate their repair‡. EMBO J. 40, e106018 (2021).

30. Liu, S. et al. RNA polymerase III is required for the repair of DNA double-strand breaks by homologous recombination. Cell 184, 1314–1329.e10 (2021).

31. Sollier, J. et al. Transcription-coupled nucleotide excision repair factors promote R-loop-induced genome instability. Mol. Cell 56, 777–785 (2014).

32. Tsherniak, A. et al. Defining a Cancer Dependency Map. Cell 170, 564–576.e16 (2017).

33. Inoue, K. & Fry, E. A. Haploinsufficient tumor suppressor genes. Adv Med Biol 118, 83– 122 (2017).

34. Solimini, N. L. et al. Recurrent hemizygous deletions in cancers may optimize proliferative potential. Science 337, 104–109 (2012).

35. Schachter, N. F. et al. Single allele loss-of-function mutations select and sculpt conditional cooperative networks in breast cancer. Nat. Commun. 12, 5238 (2021).

36. Vidotto, T. et al. Pan-cancer genomic analysis shows hemizygous PTEN loss tumors are associated with immune evasion and poor outcome. Sci. Rep. 13, 5049 (2023).

37. Malcikova, J. et al. Monoallelic and biallelic inactivation of TP53 gene in chronic lymphocytic leukemia: selection, impact on survival, and response to DNA damage. Blood 114, 5307–5314 (2009).

38. Su, S.-G. et al. An E2F1/DDX11/EZH2 Positive Feedback Loop Promotes Cell Proliferation in Hepatocellular Carcinoma. Front. Oncol. 10, 593293 (2020).

39. Dhar, S., Datta, A. & Brosh, R. M., Jr. DNA helicases and their roles in cancer. DNA Repair 96, 102994 (2020).

40. Yau, E. H. et al. Genome-Wide CRISPR Screen for Essential Cell Growth Mediators in Mutant KRAS Colorectal Cancers. Cancer Res. 77, 6330–6339 (2017).

41. Hart, T. et al. High-Resolution CRISPR Screens Reveal Fitness Genes and Genotype-Specific Cancer Liabilities. Cell 163, 1515–1526 (2015).

42. Han, K. et al. CRISPR screens in cancer spheroids identify 3D growth-specific vulnerabilities. Nature 580, 136–141 (2020).

43. Ashworth, A. & Bernards, R. Using functional genetics to understand breast cancer biology. Cold Spring Harb. Perspect. Biol. 2, a003327 (2010).

44. Chen, J. J., Silver, D., Cantor, S., Livingston, D. M. & Scully, R. BRCA1, BRCA2, and Rad51 operate in a common DNA damage response pathway. Cancer Res. 59, 1752s– 1756s (1999).

45. Prakash, R., Zhang, Y., Feng, W. & Jasin, M. Homologous recombination and human health: the roles of BRCA1, BRCA2, and associated proteins. Cold Spring Harb. Perspect. Biol. 7, a016600 (2015).

46. Hirose, T. et al. A spliceosomal intron binding protein, IBP160, links position-dependent assembly of intron-encoded box C/D snoRNP to pre-mRNA splicing. Mol. Cell 23, 673– 684 (2006).

47. Chang, L., Ruiz, P., Ito, T. & Sellers, W. R. Targeting pan-essential genes in cancer: Challenges and opportunities. Cancer Cell 39, 466–479 (2021).

48. da Costa, A. A. B. A., Chowdhury, D., Shapiro, G. I., D’Andrea, A. D. & Konstantinopoulos, P. A. Targeting replication stress in cancer therapy. Nat. Rev. Drug Discov. 22, 38–58 (2023).

49. Menzel, T. et al. A genetic screen identifies BRCA2 and PALB2 as key regulators of G2 checkpoint maintenance. EMBO Rep. 12, 705–712 (2011).

50. Zhang, X. D. et al. The use of SSMD-based false discovery and false nondiscovery rates in genome-scale RNAi screens. J. Biomol. Screen. 15, 1123–1131 (2010).

51. Lonowski, L. A. et al. Genome editing using FACS enrichment of nuclease-expressing cells and indel detection by amplicon analysis. Nat. Protoc. 12, 581–603 (2017).

52. Gerhauser, C. et al. Molecular Evolution of Early-Onset Prostate Cancer Identifies Molecular Risk Markers and Clinical Trajectories. Cancer Cell 34, 996–1011.e8 (2018).

53. Cerritelli, S. M. et al. Failure to produce mitochondrial DNA results in embryonic lethality in Rnaseh1 null mice. Mol. Cell 11, 807–815 (2003).

54. Li, Y. et al. Patterns of somatic structural variation in human cancer genomes. Nature 578, 112–121 (2020).

55. Carstens, C. J. Proof of uniform sampling of binary matrices with fixed row sums and column sums for the fast Curveball algorithm. Phys. Rev. E Stat. Nonlin. Soft Matter Phys. 91, 042812 (2015).

