## Supplemental figures for "Enzyme family-centred approach identifies helicases as recurrent hemizygous tumour suppressor genes"

### Extended data

#### Extended Data Figure 1

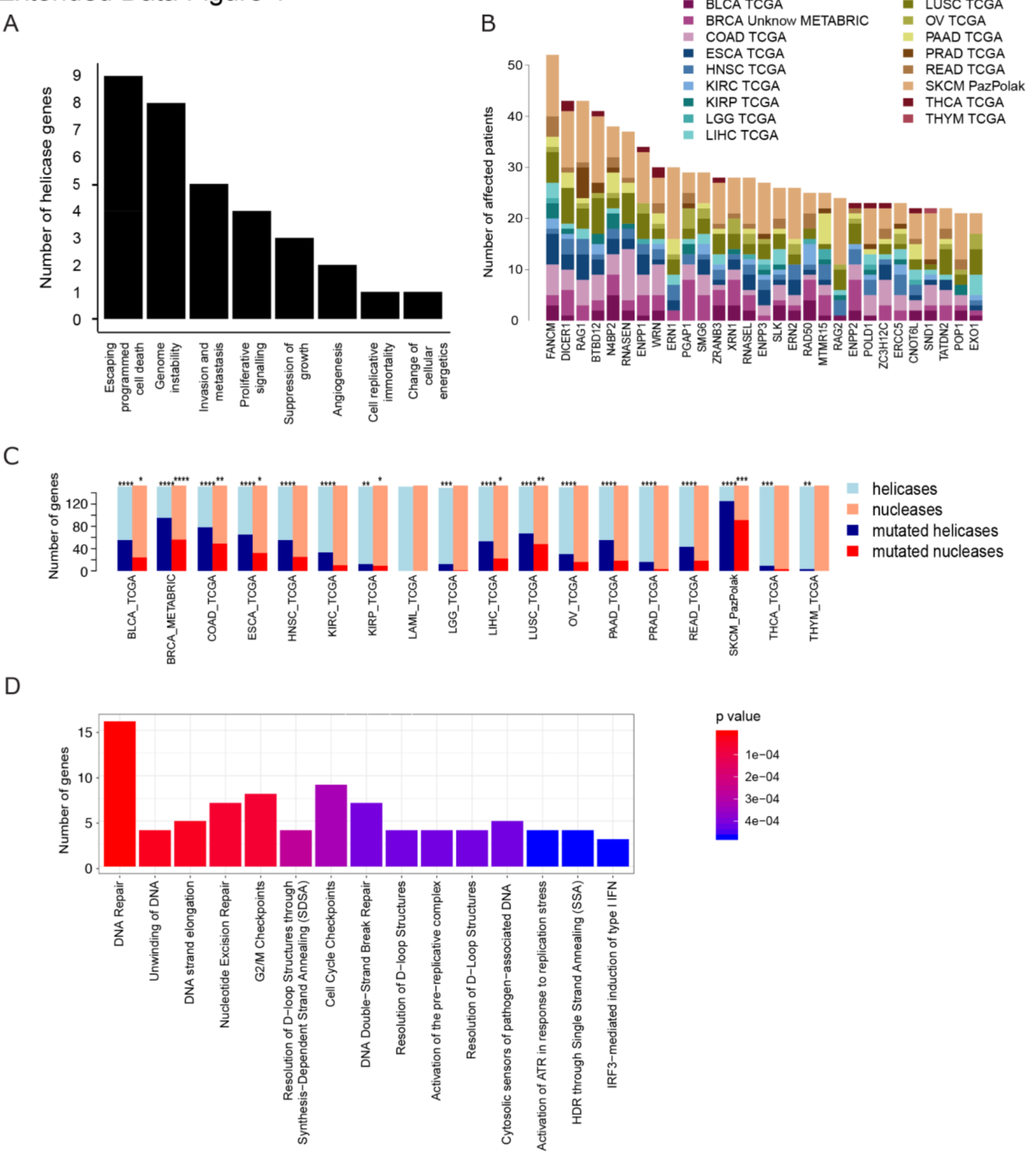


Extended Data Fig. 1

A) Hallmarks of Cancer associated with helicases in CGC.

B) Occurrence of somatic variants in nucleases: the 30 most frequently mutated nuclease genes ordered by occurrence. Each column corresponds to a gene, the colours in each column identify different tumour types, the height of a column corresponds to the number of patients in the 17 tumour-type studies (legend) carrying a mutation in the corresponding gene. See Supplementary Table 1 for the full list.

C) Number of helicase and nuclease genes mutated in 18 different tumour-type studies from TCGA and METABRIC. Asterisks denote significance level (*, $P<0.05$; **, $P<0.01$; *** $P<0.001$, **** $P<0.0001$ Fisher's exact test)

D) Pathway analysis of mutated helicase and nuclease genes in breast cancer (BRCA METABRIC study). Pathways sorted by level of p-value significance (y-axis). X-axis represents the number of genes in each pathway and the colour scale represents the adjusted p-value significance level.

#### Extended Data Figure 2

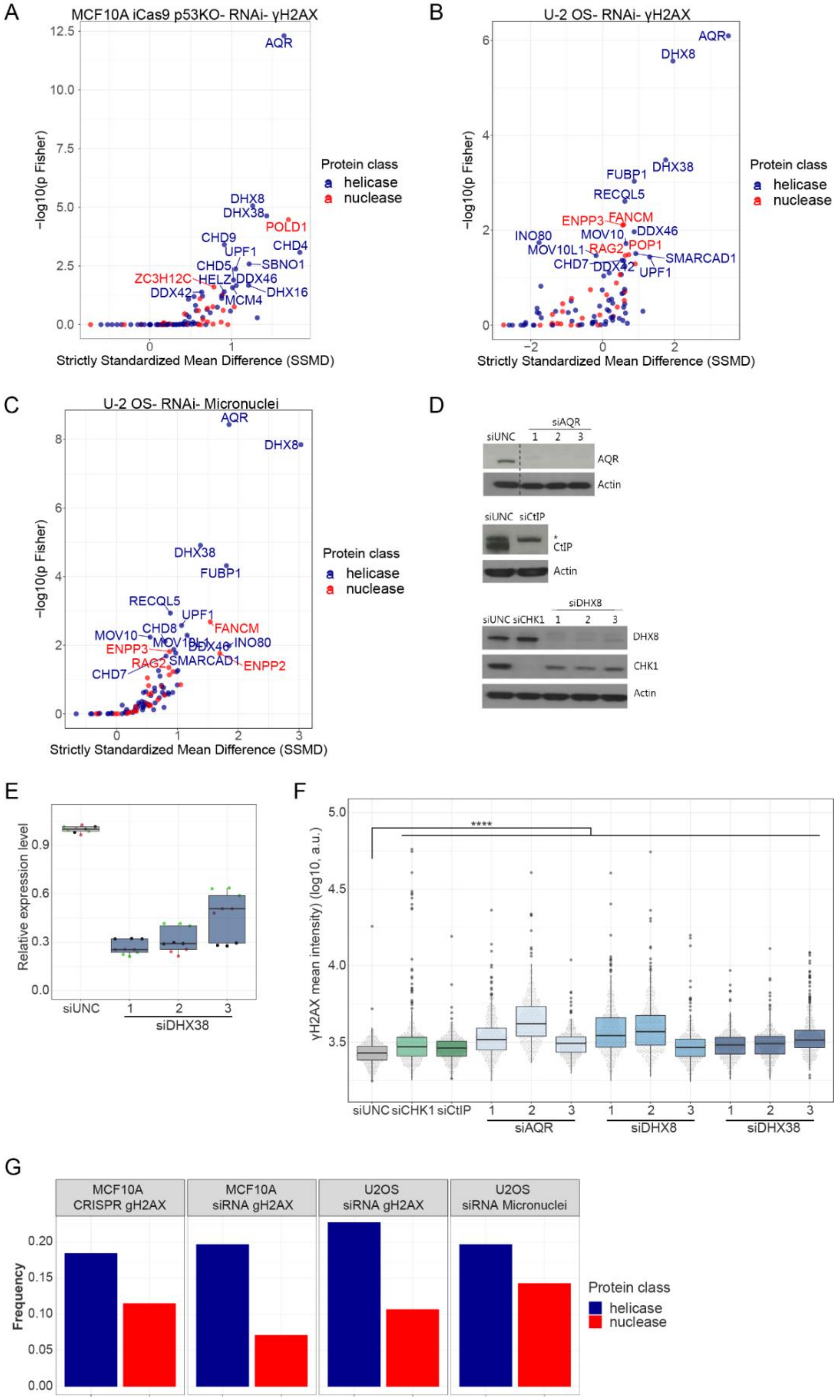

#### Extended Data Fig. 2

A) MCF10A iCas9 p53KO siRNA screen results, performed as shown in Figure 2A. On the x-axis the effect size is plotted as the strictly standardised mean difference (SSMD) in the frequency of γH2AX positive cells. The y-axis corresponds to the Fisher combined p-value from three biological replicates. Coloured by enzyme family and annotated dots represent significantly scoring genes (p Fisher <0.05).
B) U-2 OS siRNA screen results, performed as shown in Figure 2A. On the x-axis the effect size is plotted as the strictly standardised mean difference (SSMD) in the frequency of γH2AX positive cells. The y-axis corresponds to the Fisher combined p-value from three biological replicates. Coloured by enzyme family and labelled dots represent significantly scoring genes (p Fisher <0.05).
C) U-2 OS siRNA screen results for micronuclei formation, performed as shown in Figure 2A. On the x-axis the effect size is plotted as the strictly standardised mean difference (SSMD) in the frequency of micronuclei-positive cells. The y-axis corresponds to the Fisher combined p-value from three biological replicates. Coloured by enzyme family and annotated dots represent significantly scoring genes (p Fisher <0.05).
D) Immunoblot of samples in E-F and Figure 2F, assessing depletion of AQR, CtIP, CHK1 and DHX8, Actin served as a loading control. The dotted line indicates a non-continuous WB and an unspecific band detected by the CtIP antibody is labelled with an asterisk.
E) qPCR analysis of DHX38 knockdown of samples shown in F and Figure 2F. Point colours indicate samples belonging to the same biological replicate.
F) siRNA screen validation of AQR, DHX8 and DHX38-depleted U-2 OS cells. Timing as described in Figure 2A, depicting the γH2AX mean intensity. Representation of one out of three biological replicates, per sample n=450, n=150 per technical replicate. **** *P*<0.0001.
G) Analysis of the frequency of either helicase or nuclease genes scoring in each screen, normalised to the total number of helicase or nuclease genes in the target library. **** *P*<0.0001, *** *P*<0.001, * *P*<0.05, ns= not significant.

Extended Data Figure 3

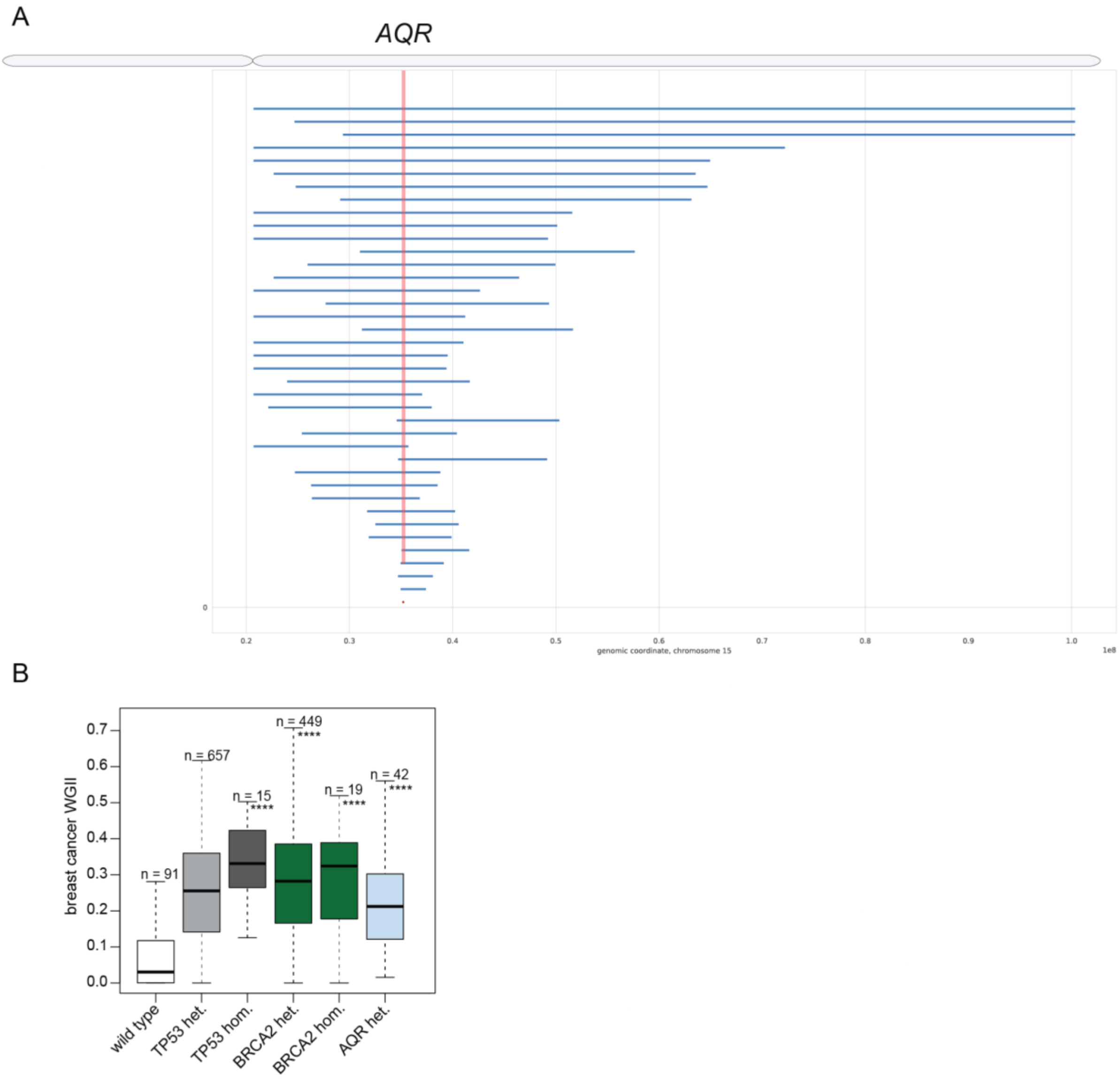


Extended Data Fig. 3

A) AQR tornado plot showing copy number alterations affecting AQR in breast cancer (PCAWG). Losses are displayed in blue and gains in red (none present).

B) Breast cancer analysis using TCGA breast cancer data of WGII for tumours with heterozygous or homozygous loss of *TP53*, *BRCA2* and *AQR*, considering samples without potential co-occurring alterations. No cancer samples had *AQR* homozygous loss (*P*=0.016, Fisher's exact test).

#### Extended Data Figure 4

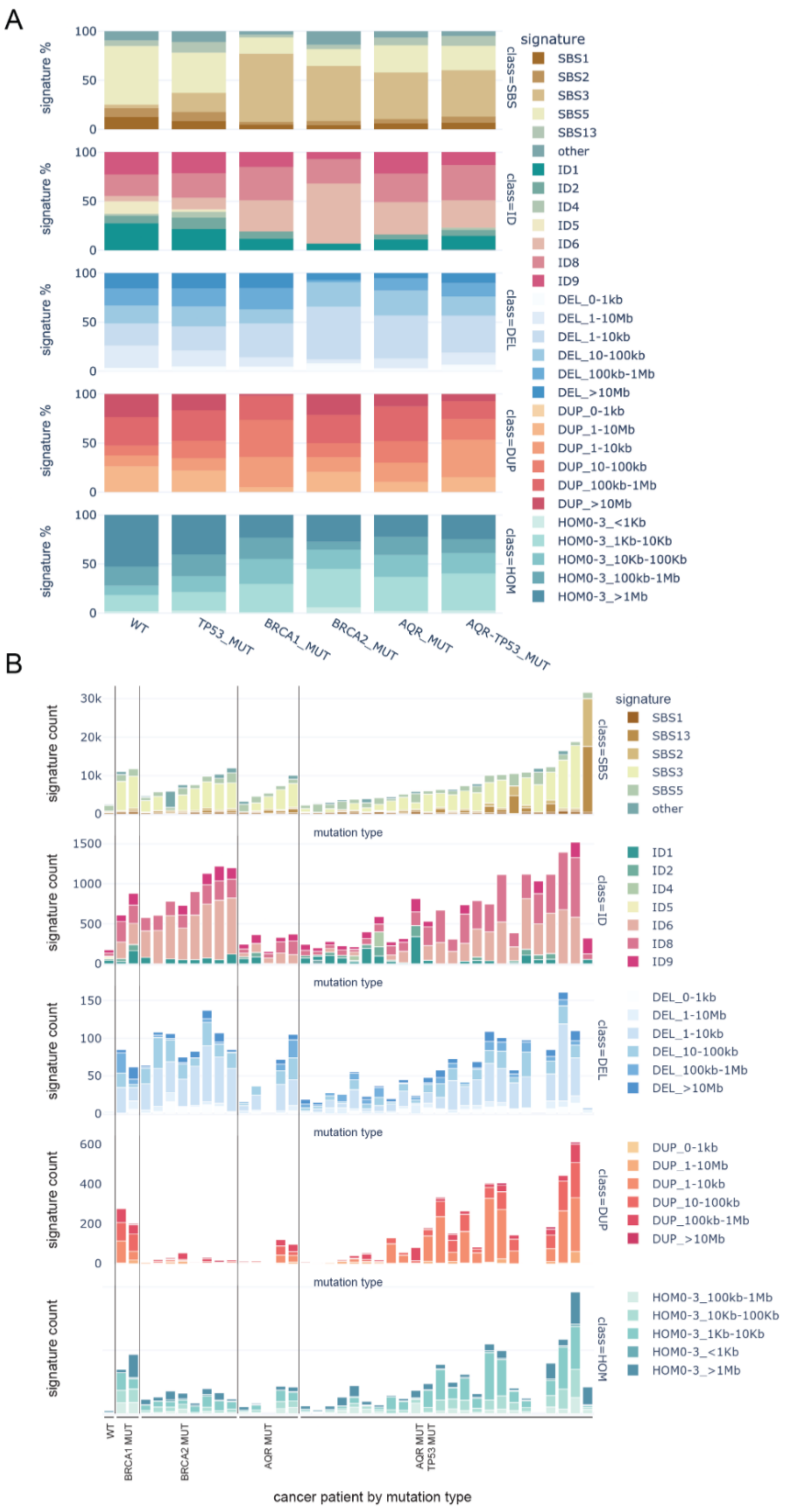

#### Extended Data Fig. 4

A) Cumulative distribution of mutational signatures for breast cancer samples with *TP53* homozygous loss (TP53 MUT), *BRCA1* homozygous loss (BRCA1 MUT), *BRCA2* homozygous loss (BRCA2 MUT), *AQR* heterozygous loss (AQR MUT) or both *AQR* heterozygous loss and *TP53* homozygous loss (AQR-TP53 MUT) or none of these mutations (WT). Top row: SBS signatures, 2nd row: ID signature, 3rd row: copy number deletion size, 4th row: copy number duplication size, 5th row: short microhomologies at different copy number sizes.

Only signatures present in at least 2% of the samples are shown.

B) Same as A) but showing the numerical signature count for the different mutational profiles. Median of all the WT samples is shown for comparison.

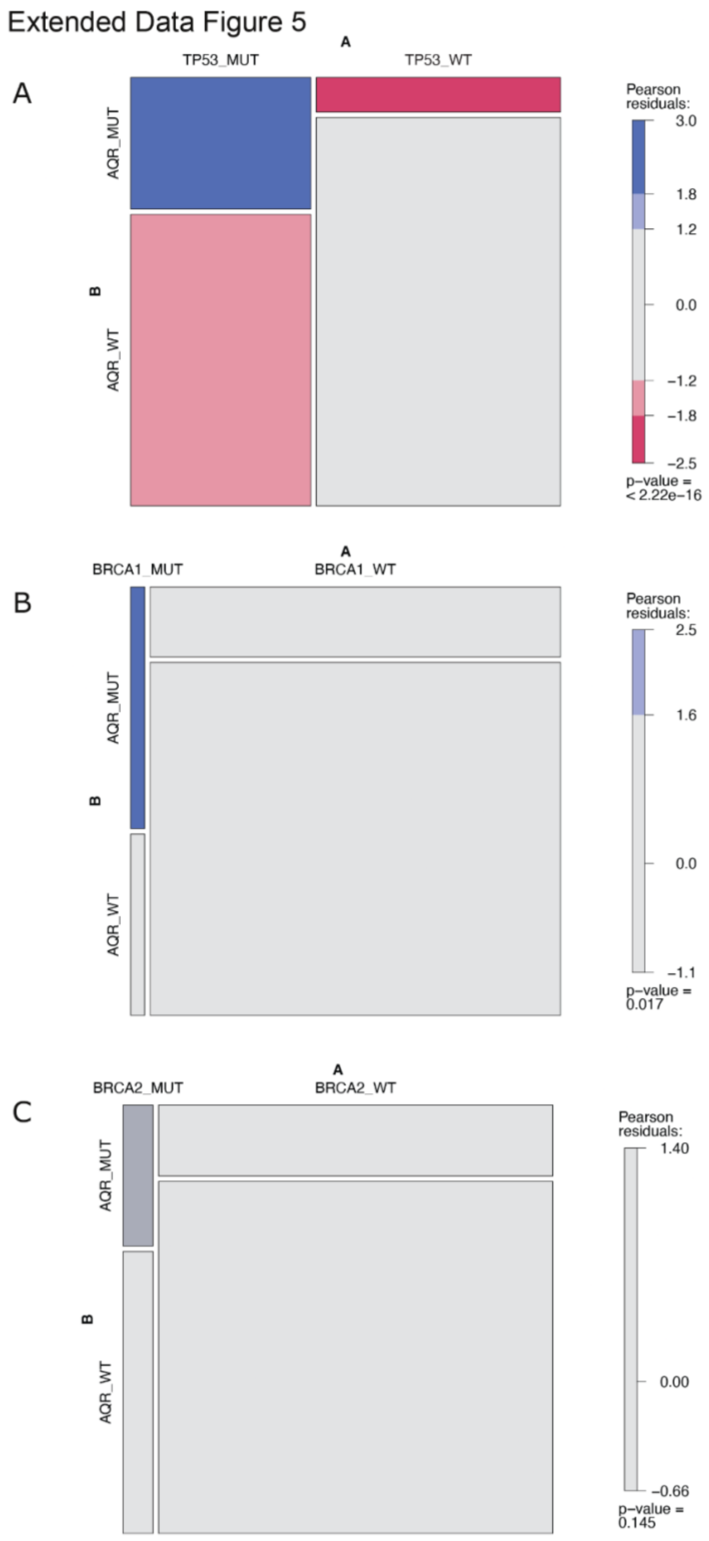


Extended Data Fig. 5

A) Co-occurrence of *AQR* and *TP53* for breast cancer samples. Contingency table of breast cancer samples with heterozygous AQR loss (AQR MUT) and/or homozygous BRCA1 loss (BRCA1 MUT). The upper left square represents cancers with co-occurring mutations and the bottom right represents

breast cancers without mutations in both enzymes, the top right are cancers exclusively with *AQR* heterozygous loss and the bottom left is cancers exclusively with *TP53* homozygous loss.
B) Similar to A) but co-occurrence of heterozygous *AQR* loss (AQR MUT) and homozygous *BRCA1* loss (BRCA1 MUT) for breast cancer samples.
C) Similar to A) but co-occurrence of heterozygous *AQR* loss (AQR MUT) and homozygous *BRCA2* loss (BRCA2 MUT) for breast cancer samples.
p-values for Fisher's exact is shown at the bottom right, with a colour gradient top right side relative to the Pearson residuals of the co-occurrence. Blue represents co-occurrence and red represents mutual exclusivity.

#### Extended Data Figure 6

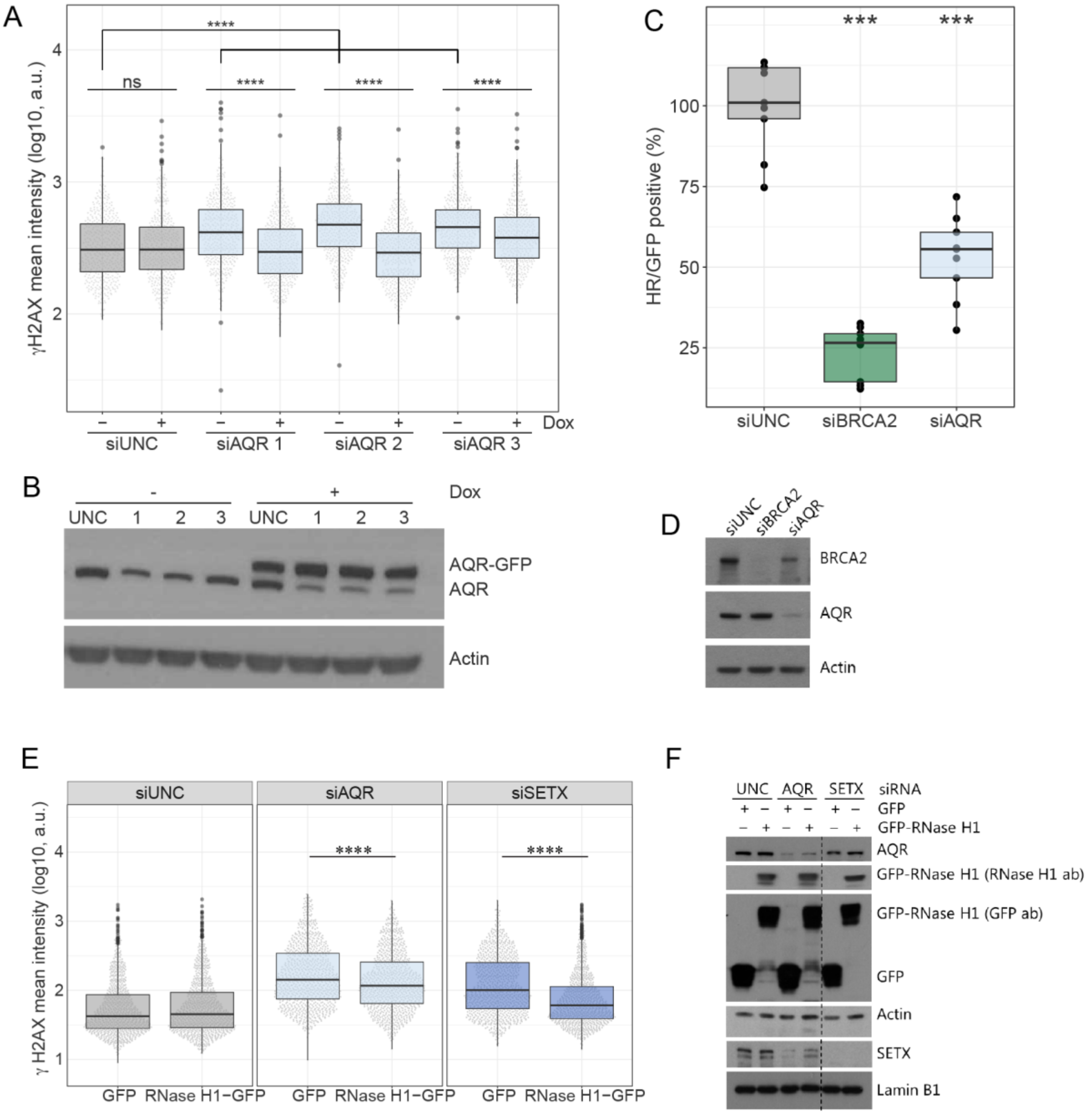


Extended Data Fig. 6

A) Analysis of AQR re-expression on genome instability in MCF10A cells with Dox-inducible AQR-GFP expression. The y-axis corresponds to the γH2AX mean intensity. The numbers 1-3 represent cells transfected with three different AQR siRNAs. Significance levels were calculated for the comparison of all three AQR siRNAs with siUNC in the absence of AQR-GFP (-Dox) as well as comparing γH2AX levels in non-induced (-Dox) vs induced (+Dox) cells for each siRNA, ****

*P*<0.0001.Representation of one out of three biological replicates, pr sample n=450, n=150 per technical replicate. **** *P*<0.0001.

B) Immunoblot of samples in B showing AQR depletion as well as AQR-GFP expression. Actin was used as a loading control.

C) Analysis of HR-efficiency in AQR- or BRCA2-depleted U-2 OS cells using the DR-GFP reporter in U-2 OS cells. 48 h post siRNA-transfection cells were transfected with I-Sce1 plasmid and harvested 24 h later. The x-axis represents the HR efficiency in S and G2 cells normalised to the siUNC control. Nine biological replicates (n=9) are represented, p-value is calculated using Wilcoxon ranksum test. ****P*<0.001.

D) Immunoblot of samples shown in D blotted using antibodies targeting AQR, BRCA2 and Actin, which was used as a loading control.

E) Analysis of γH2AX mean intensity following GFP or RNase H1-GFP expression (24 h) in AQR and SETX-depleted U-2 OS cells. γH2AX mean intensity in cells transfected with either AQR or SETX siRNA compared with siUNC control showed a significant increase in γH2AX mean intensity (p<0.0001). Representation of one out of three biological replicates, per sample n=450, n=150 per technical replicate. **** *P*<0.0001.

F) Immunoblot of samples shown in F evaluating AQR and SETX protein levels, and GFP and GFP-RNase H1 expression. Actin was used as a loading control for AQR and GFP blots, Lamin B1 was used as a loading control for the SETX blot.

### Extended Data Figure 7

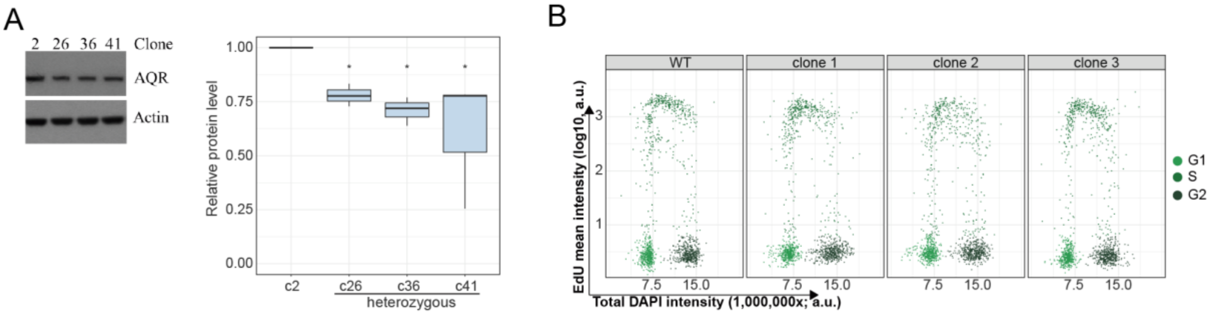


Extended Data Fig. 7

A) Analysis of AQR protein levels by WB in AQR heterozygous MCF10A iCas9 p53KO clones, immunoblot and quantification, same samples as in Figure 4B. Actin served as a loading control. Analysis of three independent WB, * P< 0.05.

B) Cell cycle analysis of wild type or three heterozygous *AQR* MCF10A iCas9 p53KO clones. The x-axis corresponds to the total DAPI intensity and the y-axis to the EdU mean intensity. Cells were gated into G1, S and G2 phase based on total DAPI and mean EdU intensity. Representation of one out of three biological replicates, n=450 per sample, n=150 per technical replicate.

#### Extended Data Figure 8

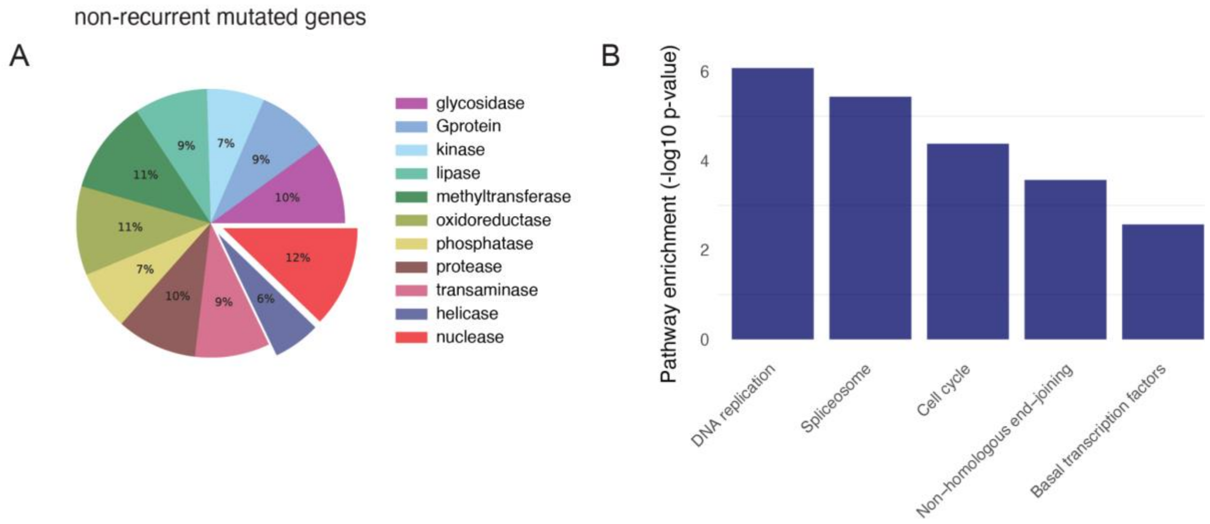


##### Extended Data Fig. 8

A) Non-recurrently mutated genes (less than 3 focal mutations, or a pan-cancer focal recurrence < 0.1%), essentiality (Dependency score < -1) and not found biallelic inactivated (<0.5%).
B) Significantly enriched KEGG pathways for Hemizygous driver genes. Only significant pathways are shown.
